# Dual-targeting strategy to repurpose Cetuximab with HFn nanoconjugates for immunotherapy of triple-negative breast cancer

**DOI:** 10.1101/2025.02.03.634255

**Authors:** Linda Barbieri, Lucia Salvioni, Andrea Banfi, Stefania Garbujo, Luisa Fiandra, Chiara Baioni, Marco Giustra, Lucia Morelli, Gianni Frascotti, Miriam Colombo, Metello Innocenti, Davide Prosperi

**Affiliations:** Department of Biotechnology and Biosciences, University of Milano-Bicocca, Piazza Della Scienza 2, 20126 Milan, Italy

**Keywords:** Triple-negative breast cancer, Cetuximab, ferritin, nanoconjugates, epidermal growth factor receptor, transferrin receptor, immunotherapy

## Abstract

Triple-negative breast cancer (TNBC) is a highly aggressive and treatment-resistant malignancy, characterized by the lack of targeted therapies and poor clinical outcomes. Here, we present a dual-targeting strategy combining the anti-EGFR monoclonal antibody Cetuximab (CTX) with H-ferritin (HFn), a nanoparticle targeting transferrin receptor 1 (TfR1), for potential immunotherapy in CTX-resistant tumors. The HFn-CTX nanoconjugate exhibited favorable biophysical properties, including a hydrodynamic size of <30 nm, and significantly enhanced antibody-dependent cellular cytotoxicity (ADCC) in TNBC spheroids compared to CTX alone. Conversely, glioblastoma spheroids did not exhibit comparable reactivity. This effect correlated with elevated cell-surface EGFR expression and plasma-membrane lingering of the nanoconjugate in TNBC cells, facilitating robust immune activation. Biodistribution studies showed selective accumulation of the HFn-CTX nanoconjugate in TNBC tumors in vivo. These findings highlight the potential of HFn-CTX nanoconjugates to repurpose CTX for refractory cancers that express EGFR at high levels like TNBC, leveraging dual-receptor targeting to amplify immune-mediated cytotoxicity and overcome resistance.

## Introduction

Breast cancer (BC) is the most frequent malignancy in women worldwide and remains the second cause of cancer death among females. BC represents about 12 percent of all new cancer cases each year and 25 percent of all cancers in women. Early-stage non-invasive or locally invasive BC can be cured in almost 70–80% of patients. In contrast, advanced (metastatic) BC remains incurable, because the available therapeutic options can only delay progression or serve as palliative treatments.^1^

The breast tumors that do not express estrogen receptor, progesterone receptor, or human epidermal growth factor receptor 2 (HER2, aka ErbB-2) are referred to as triple-negative breast cancer (TNBC), which accounts for approximately 11-20 percent of all female BC cases and has the highest mortality rate.^2^ In keeping with this, TNBC shows a high histological grade, a high proliferation rate, a high risk of relapse, short progression-free and overall survival.^3,4^

TNBC treatment is challenging due to the lack of obvious drug targets, with surgery, Hence, developing effective targeted therapies for TNBC is an urgent medical need. In this regard, it is tantalizing that 40-60% of all TNBC overexpress epidermal growth factor receptor 1 (EGFR, aka HER1, aka ErbB-1), a well-known proto-oncogene.^7,8^ Several EGFR-targeting mAbs have been developed in recent years. Among them, Cetuximab (CTX) binds to the extracellular domain of EGFR with high affinity and competes with endogenous ligands thereby inhibiting EGFR signaling. This produces anti-tumor effects such as cell-cycle arrest, induction of apoptosis, inhibition of angiogenesis and metastasis, and enhanced sensitivity to radio- or chemo-therapy.^9^ Besides anticancer effects mediated by the binding to cell-surface EGFR, CTX can promote immune cell-induced tumor killing through a process called antibody-dependent cellular cytotoxicity (ADCC). The ADCC induced by CTX is EGFR-dependent and mediated by the activation of natural killer (NK) cells. Notably, these immunomodulatory properties were found to contribute to the anti-tumor effects observed in the clinic.^10,11^ Yet, the use of CTX as an anti-EGFR therapeutic has only been approved for the treatment of patients with wild-type *RAS* metastatic colorectal cancer and head and neck squamous cell cancer.

This is certainly related to the general limitations of many mAbs, which include inadequate pharmacokinetics, short circulation half-life, low-tumor penetration, inability to cross biological barriers, and several types of resistance mechanisms.^12–14^ Nanotechnology can help bypass some of these drawbacks in cancer therapy, and different types of EGFR-targeted nanoparticles relying on mAbs have shown some effects on TNBC *in vitro*.^15–17^ Although mAbs-based active tumor targeting makes these nanoparticles potentially superior to those counting on the enhanced permeability and retention (EPR) effect for accumulation in the tumor, selectivity remains an issue. Indeed, they would be retained in all bodily tissues expressing the EGFR at appreciable levels.^18^

An elegant approach to enhance the specificity and levels of nanoparticle accumulation within tumors would be combinatorial targeting of tumor surface antigens. A prime example is transferrin receptor 1 (TfR1, aka CD71), a cell-surface antigen overexpressed in 98% of human solid cancers, including TNBC.^19–22^ For this reason, the high-affinity TfR1 ligand H-ferritin (HFn), a fully biocompatible recombinant round-shaped cage protein that can be loaded with various payloads, has attracted much interest in the nanomedicine field. As the HFn nanoparticles exhibited good tumor targeting and the ability to cross the blood brain barrier,^23–26^ HFn has been exploited as a drug delivery system for chemotherapeutics improving their accumulation in the tumor while reducing adverse side effects.^27^ Moreover, the surface of the HFn cage can be modified, both genetically and chemically, to add functional moieties. In particular, surface primary amines were successfully conjugated to peptides, antibodies, or fluorophores.^28,29^

Here, we provide a proof of principle of combinatorial tumor targeting that leverages both the high affinity and specificity of HFn and CTX for their cognate receptor to develop a novel HFn-CTX nanoconjugate and repurpose CTX for the treatment of a wider array of cancers. We set up a tailored pipeline to produce and purify HFn-CTX nanoconjugate, the uptake of which was assessed in TNBC and glioblastoma (GBM) cells expressing high EGFR levels and harboring K-*ras* or *PTEN* mutations associated with CTX resistance.^30,31^ To weigh the HFn-CTX nanoconjugate as a valuable tool to kill CTX-resistant cells, we measured the immune-mediated anti-cancer activity of the nanoconjugate in 3D cultures. Surprisingly, we discovered that its ability to induce ADCC was correlated with the cell-surface EGFR and TfR1 levels and internalization kinetics. Furthermore, the HFn-CTX nanoconjugate showed a favorable biodistribution and improved tumor accumulation and retention in a preclinical mouse model of TNBC. Taken together, these results open the door to repurposing CTX with HFn nanoconjugates for immunotherapy of patients with TNBC or other cancers ineligible to standard CTX treatments.

## Results

### Production and biophysical characterization of CTX-conjugated HFn nanoparticles

To produce the HFn-CTX nanoconjugate, we initially employed a standard protocol for mAbs-conjugated HFn nanoparticles (NPs).^24,32^ In brief, HFn nanoparticles were produced and purified using established methods,^33^ whereas CTX was reacted with a 5 kDa PEG-based heterobifunctional crosslinker harbouring one *N*-hydroxysuccinimidyl ester (NHS) and one maleimide (Mal) group. Next, the resulting PEGylated antibody was conjugated with the HFn nanoparticles to synthesize the HFn-CTX nanoconjugate (Fig. 1A). We optimized the stoichiometries of the various reactants, exploiting those allowing the production of other HFn-mAbs nanoconjugates as a reference point.^24^ For the HFn-CTX nanoparticles, 1:1 was the molar ratio between the two protein species that we selected to ensure that both HFn and CTX would be adequately spaced to bind their cognate receptors and to minimize accumulation in the liver.^34^ Under these conditions, batches of HFn-CTX nanoparticles with reproducible physicochemical properties could be synthesized (Table S1).

**Figure 1.**
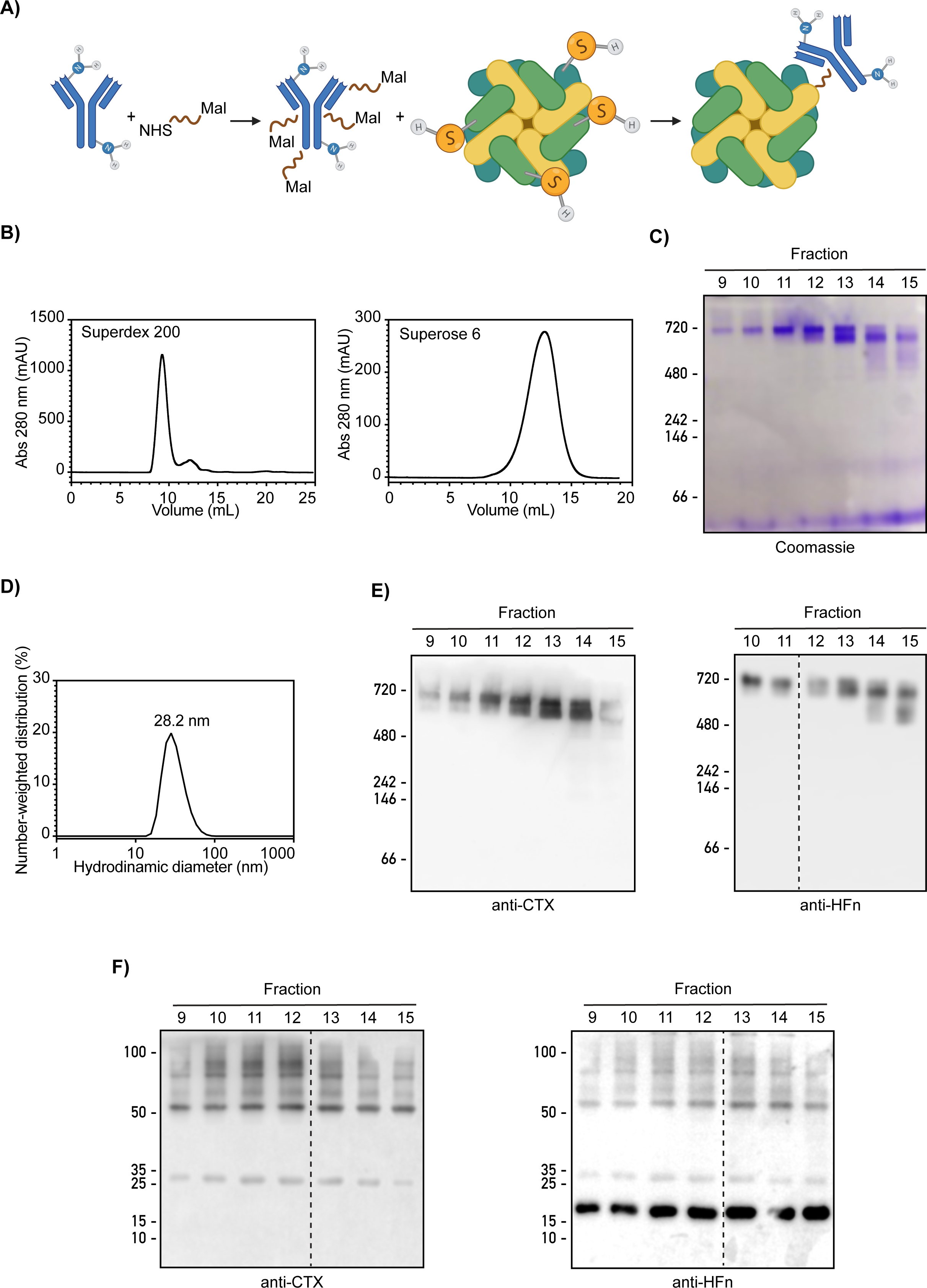
Synthesis, purification, and characterization of the HFn-CTX nanoconjugate. **(A)** Schematic representation of the two reactions leading to the synthesis of the HFn-CTX nanoconjugate. PEGylation of the mAb is followed by conjugation with HFn nanoparticles. **(B)** Representative elution profiles of the purification of the HFn-CTX nanoconjugate by SEC-FPLC. (*Left*) Elution profile of the products obtained from the synthetic reaction (A) on Superdex 200. (*Right*) Elution of the void fractions (9-11) from the Superdex 200 column loaded on a Superose 6. **(C** and **E)** Characterization of the native HFn-CTX nanoconjugate. Protein molecular weight markers, purified HFn and CTX standards, and equal volumes (10 µL) from the indicated fractions (fr) of the HFn-CTX complex eluted from the Superose 6 column were separated by native page electrophoresis (running gel: 6%) and blotted with the indicated antibodies. **(D)** Size of the HFn-CTX nanoconjugate. DLS was used to determine the hydrodynamic size of the purified HFn-CTX nanoconjugate. **(F)** Characterization of the denatured HFn-CTX nanoconjugate. The fractions analyzed in (C) were separated by SDS-PAGE (running gel: 10%) and blotted as indicated. Dashed lines indicate removal of intervening lanes.

We compared different HFn *vs*. Alexa Fluor 488 (AF488) dye molar ratios, as well as different protein concentrations and reaction conditions, to optimize protein labeling (Table S2). Based on the degree of labeling (DOL), we chose a 1:50 molar ratio and a HFn concentration of 5 mg mL^−1^. Similar experiments were conducted to optimize the labeling of CTX, using the AF647 dye (Table S3). PEG was also added to the reaction because this linker competes with AF647 for the binding to the amine groups of CTX. We selected a 1:10 molar ratio and a CTX concentration of 5 mg mL^−1^ (Table S3) to avoid over-labeling, which could cause both aggregation and reduce the specificity of the antibody. This streamlined the production of different variants of fluorescent nanoconjugates, among which we characterized *in vitro* and *in vivo* those labeled on HFn and CTX with AF647 and AF750 CTX, respectively, for their advantageous emission spectra (Table S4 and S5).

The nanoparticles were purified by size exclusion chromatography (SEC) and subsequently analysed by SDS-PAGE and western blot (Fig. 1B, C, D and E). We initially took advantage of the SEC setup utilized for HFn-Trastuzumab nanoconjugates,^24^ but we were unable to identify distinct peaks or fractions rich in the HFn-CTX nanoconjugate while depleted of each unreacted species (Fig. S1A). Hence, we established a new two-step purification protocol, which is described in Fig. S1B. This allowed us to retrieve good amounts of highly pure HFn-CTX nanoconjugates (Fig. 1C, E, and F). The hydrodynamic size of this nanoconjugate was lower than 30 nm, as measured by dynamic light scattering (DLS) (Fig. 1D).

### The HFn-CTX nanoconjugate boosts ADCC in TNBC spheroids

To evaluate the ADCC induced by the HFn-CTX nanoconjugate, we quantified apoptosis induced by human peripheral blood mononuclear cells (PBMCs), which contain NK cells. Upon activation, NK cells polarize and exocytose Perforin and Granzyme B granules, which induce apoptotic cell death by causing the cleavage of procaspase 8/10 and the ensuing activation of caspase 3/7.^35^

U-87 MG and MDA-MB-231 spheroids were pre-incubated with either CTX, or HFn-CTX, or HFn prior to adding fresh IL2-activated PBMCs. The spheroids were lysed 24 and 48 hours later, and caspase 3/7 activity was measured. The TNBC spheroids treated with the nanoconjugate showed a more prominent caspase activation than those treated with either CTX or HFn alone (Fig. 2A, *right*). Unexpectedly, the ADCC was much milder in the GBM spheroids, and only the CTX-treated ones exhibited a mild yet significant response (Fig. 2A, *left*). Importantly, the enhanced ADCC observed in the TNBC spheroids was independent of size (Fig. 2B) or penetration of the HFn-CTX nanoconjugate, as shown by automated image analysis of cryosections obtained from parallel GBM and TNBC spheroids (Fig. S2). The HFn-CTX nanoconjugate was mainly detected in the outer layers of the spheroids, whereas the HFn nanoparticles were slightly more infiltrated, particularly in the MDA-MB-231 spheroids (Fig. S2A and B). The detected signals were specific, as shown by imaging untreated spheroids (Fig. S2C). Of note, accumulation and distribution within spheroids did not seemingly change by increasing the incubation time from 30 minutes to 4 h (Fig. S2 A and B). This is consistent with both the receptor-mediated uptake of HFn and CTX further enhancing high affinity binding of HFn nanoparticles on the cell surface.

**Figure 2.**
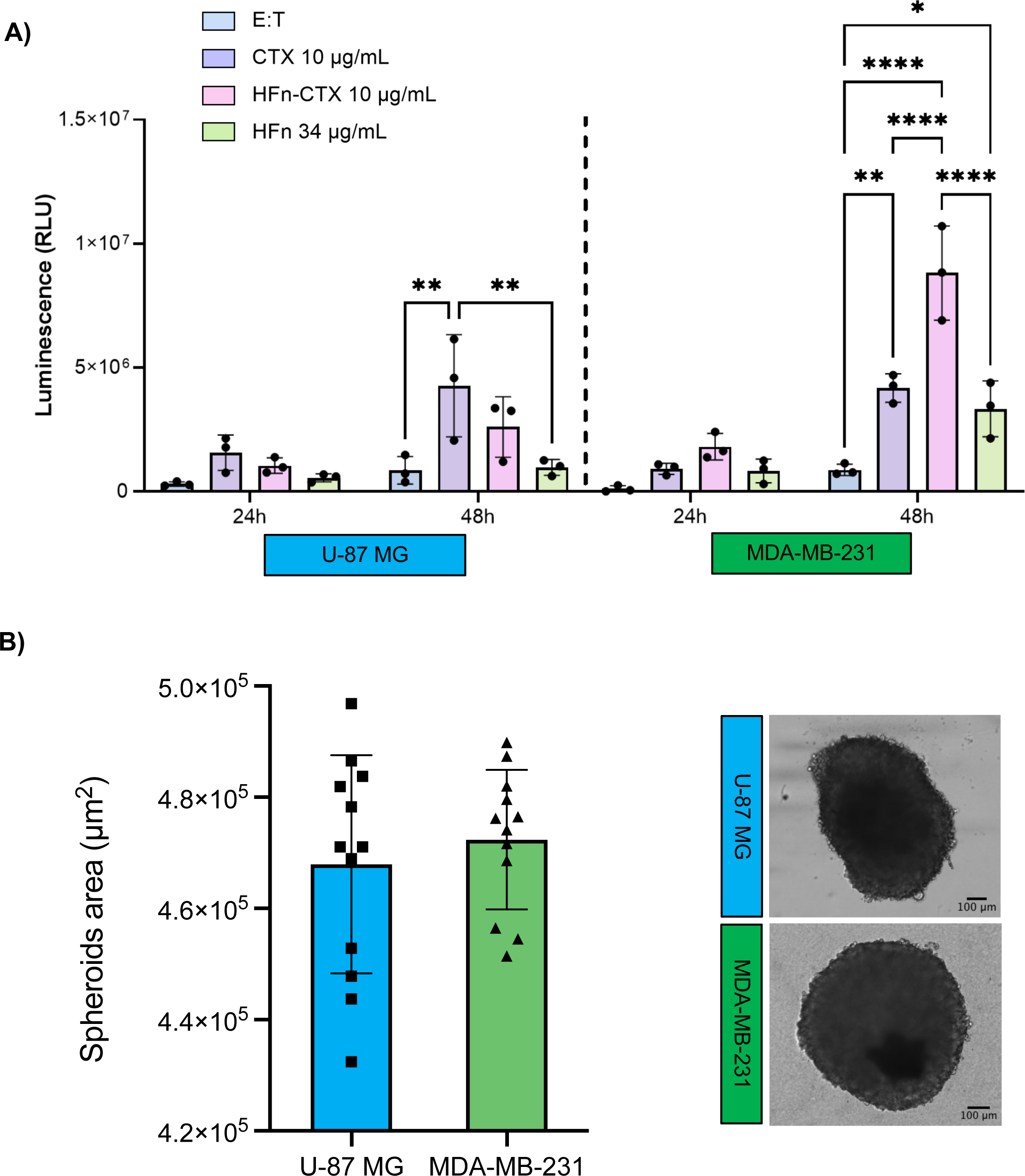
The HFn-CTX nanoconjugate promotes ADCC in TNBC but not GBM spheroids. **(A)** Caspase activity in U-87 MG and MDA-MB-231 spheroids treated with of CTX (10 μg mL^−1^), or HFn-CTX (10 μg mL^−1^), or HFn (34 μg mL^−1^), was measured at 24 and 48 h upon addition of IL-2-activated PBMCs. Bar graphs show the mean ± s.e. of 3 independent replicates (each composed of three technical replicates). * = p < 0.05; ** = p < 0.01; **** p < 0.0001 (Two-way ANOVA). **(B)** Area of U-87 MG and MDA-MB-231 spheroids. Each square or triangle represents the area of a single spheroid at 5 days of seeding. On the right, representative images of U-87 MG and MDA-MB-231 spheroids at 5 days of seeding.

### The uptake of the HFn-CTX nanoconjugate is slower in TNBC than in GBM cells

To gain insight into why the HFn-CTX nanoconjugate promoted a robust ADCC in TNBC but not in GBM spheroids, we analyzed its uptake and compared it to that of the HFn nanoparticles. The TfR1 undergoes rapid and constitutive internalization via clathrin-mediated endocytosis and then recycles back to the plasma membrane,^36,37^ whereas the EGFR can be internalized through different routes, which contribute to determining whether it is recycled or degraded.^38,39^

Time-course experiments revealed that the HFn-CTX nanoconjugate was rapidly internalized in U-87 MG cells, where it localized in punctate structures likely corresponding to the endolysosomal system, as early as at 5 min of incubation (Fig. 3A and Fig. S3). Interestingly, the nanoconjugate and the HFn nanoparticles showed similar uptake kinetics (Fig. 3A). Surprisingly, the nanoconjugate lingered on the plasma membrane of MDA-MB-231 cells much longer (up to 30 minutes), and it took longer to accumulate in intracellular compartments as compared to the HFn nanoparticles (Fig. 4A, and Fig. S3).

**Figure 3.**
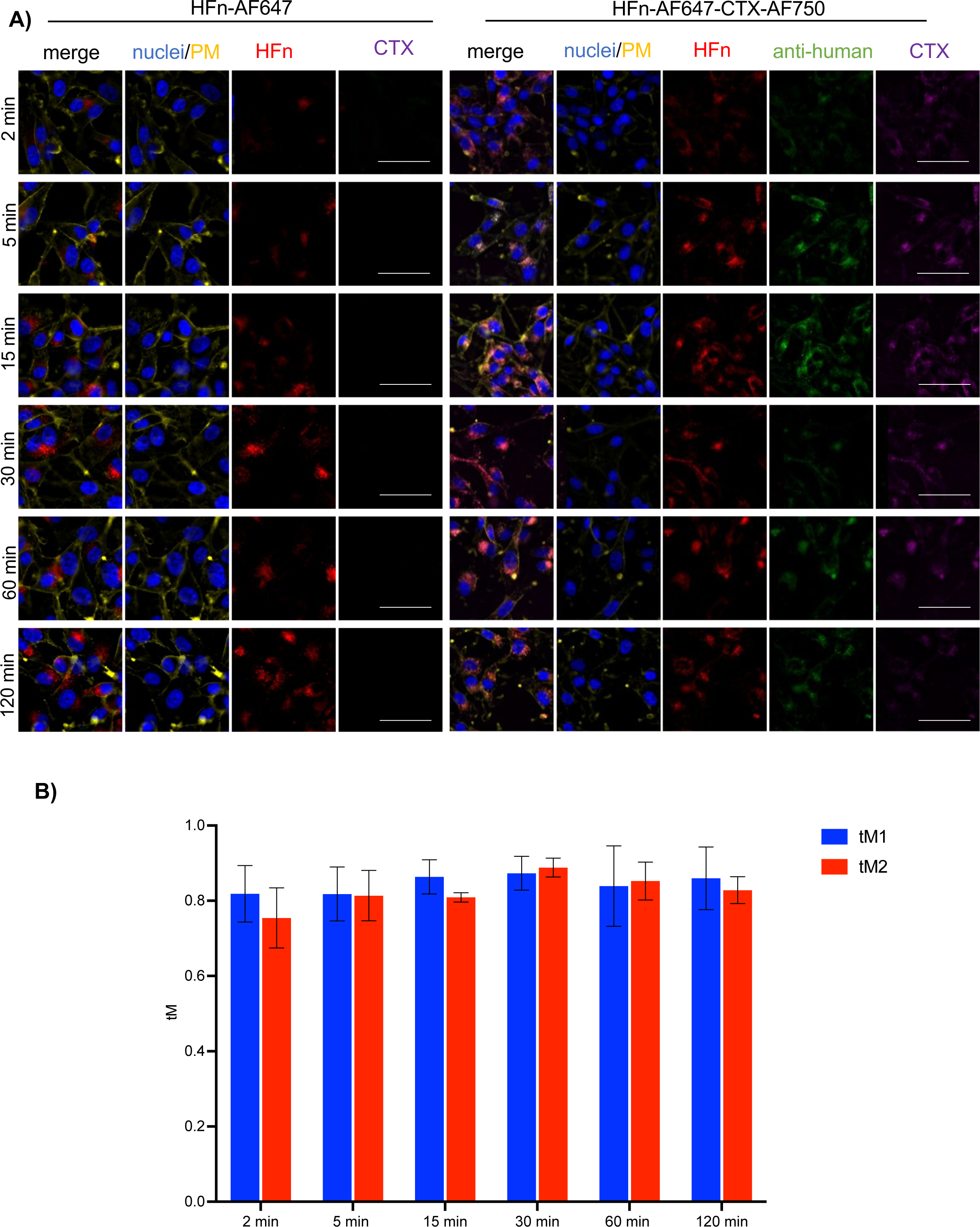
The HFn-CTX nanoconjugate and the HFn nanoparticles show similar uptake kinetics in GBM cells. **(A)** Representative images of U-87 MG cells incubated with double labelled HFn-CTX or labelled HFn (0.1 mg mL^−1^ each) for 2, 5, 25, 30, 60 and 120 min. HFn-AF647 is depicted in red, CTX in magenta, and CTX detected with anti-human-AF488 in green. Plasma membrane (PM) was stained with CD44 (yellow), nuclei were stained with Hoechst (blue). Scale bar, 100 µm. **(B)** Mander’s coefficient analysis on HFn-CTX nanoconjugate and HFn nanoparticles. Bar graph shows tM1 (CTX-AF488 signal *vs.* HFn-AF647 signal, blue bars) and tM2 (HFn-AF647 signal vs CTX-AF488 signal, red bars) at the indicated time points.

**Figure 4.**
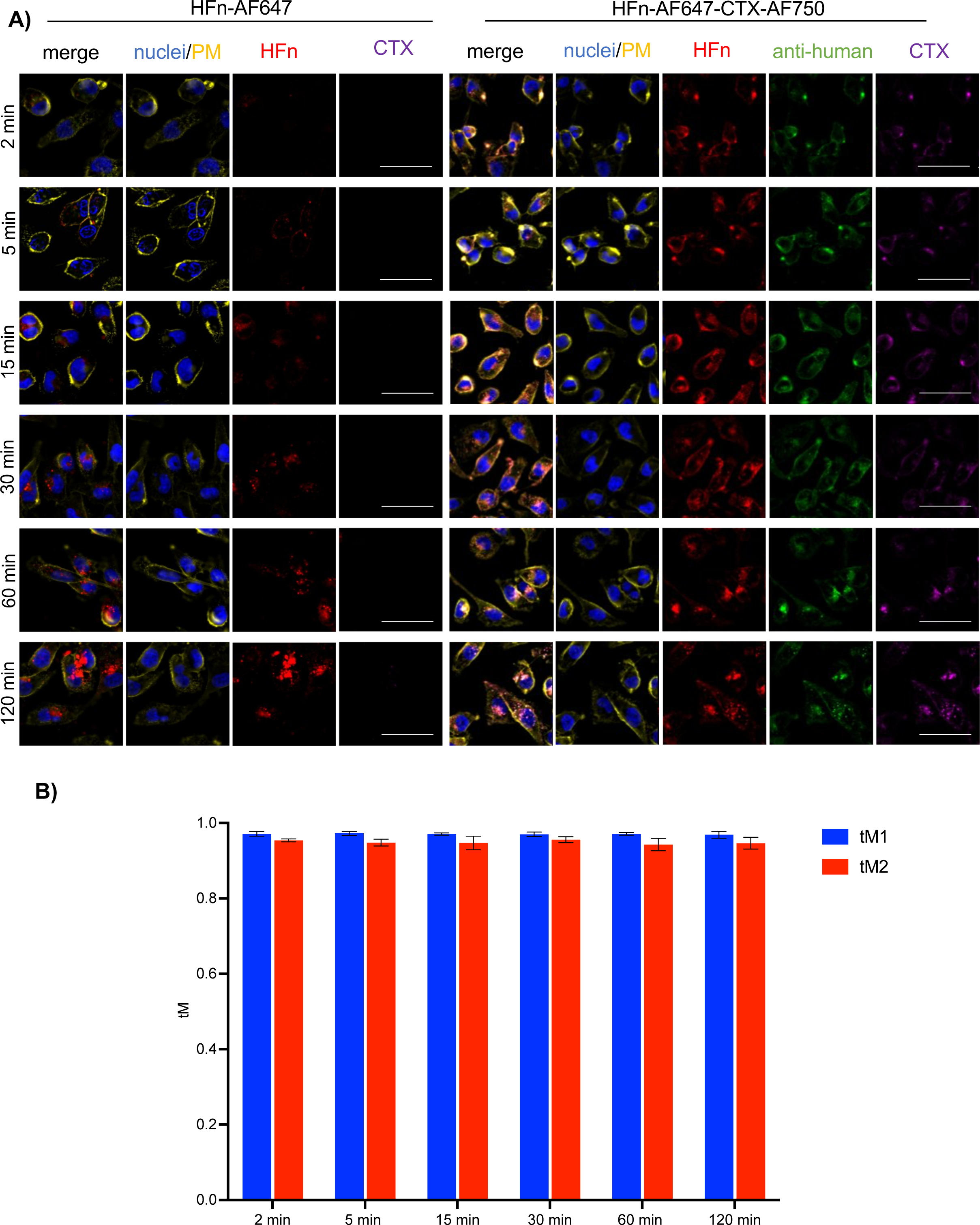
TNBC cells take up the HFn-CTX nanoconjugate more slowly than the HFn nanoparticles. **(A)** Representative images of MDA-MB-231 cells incubated with double labelled HFn-CTX or labelled HFn-647 (0.1 mg mL^−1^ each) for 2, 5, 25, 30, 60, and 120 min.. Cells were processed, stained, and imaged as in Fig. 3A. Scale bar, 100 µm **(B)** Mander’s coefficient analysis at all the tested timepoints. Data were obtained and plotted as in Fig. 3B.

The Mander’s coefficient values (tM1=CTX-AF488 signal *vs*. HFn-AF647 signal, and tM2=HFn-AF647 signal *vs* CTX-AF488 signal) were close to 1 at all the tested time points in both cell lines, indicating an almost complete colocalization between the HFn and CTX signals (Fig. 3B and 4B). These results ruled out that a cell type-specific massive dismantling of the internalized nanoconjugate could explain the above discrepancies.

Hence, the MDA-MB-231 cells seem to internalize the HFn-CTX nanoconjugate more slowly than the HFn nanoparticles, whereas both are taken up faster and with similar kinetics by the U-87 MG cells.

### TNBC and GBM cells have distinct cell-surface levels of EGFR and TfR1

Differences in the expression or cell-surface levels of TfR1 and EGFR, to which HFn and CTX bind, would impact on the internalization of the nanoconjugate and thereby account for the above observations. Thus, we sought to compare the protein levels of TfR1 and EGFR in glioblastoma and TNBC cells. Western Blot analyses showed that TfR1 was significantly more expressed in the U-87 MG than in the MDA-MB-231 cells, whereas the opposite held for the EGFR (Fig. 5A,B). Furthermore, measuring the fluorescence intensity of the HFn-CTX nanoconjugate or HFn nanoparticles at early time points (2 min) of uptake, when the degradation of internalized cargos has not yet occurred, unmasked that the signal of the bound HFn-CTX nanoconjugate was not only much higher (74x) but also increased (3x) compared to that of the HFn nanoparticles bound to the MDA-MB-231 or U-87 MG cells (Fig. 5C). These data suggested that the former cells have more binding sites for these nanoparticles.

**Figure 5.**
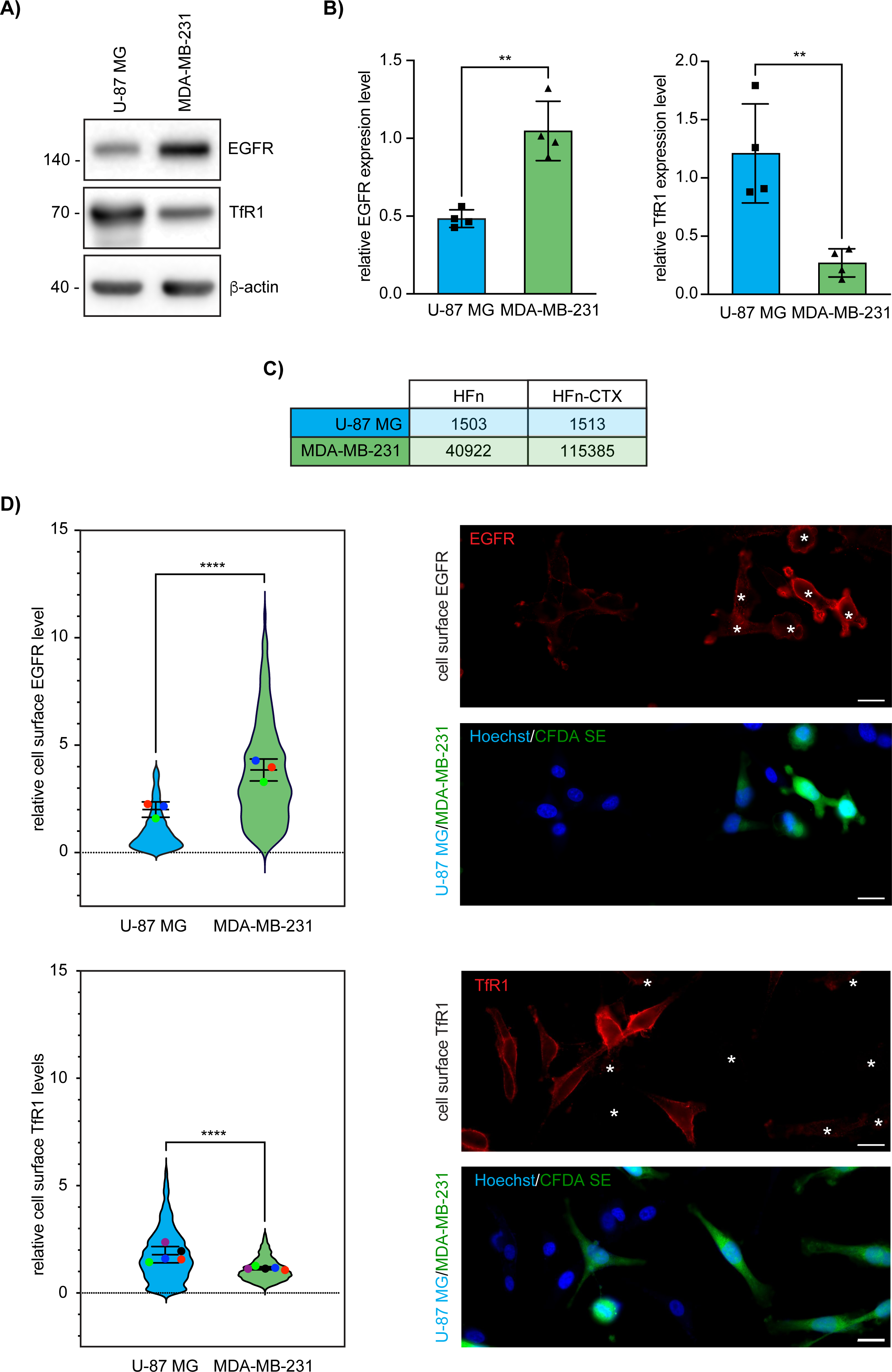
TfR1 and EGFR expression in U-87 MG and MDA-MB-231 cells. **(A)** Representative Western blot of total lysates of U-87 MG and MDA-MB-231 cells (15 µg) blotted as indicated. **(B)** Densitometric quantification of the TfR1 and EGFR signal normalized on the corresponding ß-actin signal. Bar graphs show mean ± s.e. of 3 independent replicates. ** = p < 0.01 (Two-tailed Student’s t-test). **(C)** Quantification of the fluorescence intensity of HFn alone or conjugated with CTX normalized by the number of nuclei. Data were obtained by measuring HFn-AF647 fluorescence in U-87 MG and MDA-MB-231 cells after 2 minutes of incubation with HFn or HFn-CTX. **(D)** Cell-surface EGFR and TfR1 levels as measured with CellProfiler and representative images of non-permeabilized cells stained for EGFR or TfR1. Nuclei were stained with Hoechst (blue), TfR1 and EGFR (red), CFDA SE-labeled MDA-MB-231 cells (green). White asterisks mark the position of the MDA-MB-231 cells in the anti-TfR1 and anti-EGFR micrographs. Scale bar, 20 µm. Violin superplots show the mean ± s.e. of 3 independent replicates (1665 U-87 MG cells and 1065 MDA-MB-231 cells for EGFR; 2178 U-87 MG cells and 1751 MDA-MB-231 cells for TfR1). ** = p < 0.01; **** p < 0.0001 (unpaired two-tailed T-test).

To follow up on this, we measured the relative amount of cell-surface EGFR and TfR1 in MDA-MB-231 and U-87 MG co-cultures by staining non-permeabilized fixed specimens with antibodies targeting the ectodomain of either receptor. Automated quantitative analysis of the resulting microscopy images revealed that MDA-MB-231 cells had more EGFR on the cell surface than U-87 MG cells. Conversely, cell-surface TfR1 showed the opposite trend, although differences were slightly less marked (Fig. 5D and Fig. S4).

### Biodistribution of the HFn-CTX nanoconjugate and tumor-targeting

Prompted by the sum of above results, we sought to assess the biodistribution of the HFn-CTX nanoconjugate and its ability to target TNBC tumors. MDA-MB-231 cells stably expressing a luciferase reporter were injected subcutaneously into female NOD/Scid mice. When tumors reached a volume of 100 mm^3^, typically 3 weeks post-injection, the AF647-labelled HFn-CTX nanoconjugate (5 mg kg^−1^) was injected into the tail vein, whereas PBS was used as a control. To evaluate biodistribution, healthy tumour-bearing mice were administered the HFn-CTX nanoconjugate, and epifluorescence (Epf) images were taken at 30 minutes, 1 h, 3 h, and 24 h post-injection, keeping the animals at 37 °C. After completing the imaging, the mice were sacrificed; organs and tumors were dissected and subsequently imaged at room temperature. The signal of the HFn-CTX nanoconjugate was intense in the liver, in line with the known biodistribution of HFn nanoparticles.^40^ For this reason, it was masked while whole animals were imaged (Fig. 6A). Resected tumors showed a progressive accumulation of the nanoconjugate from 30 minutes to 3 hours, when the signal peaked, followed by a marked decrease at 24 h post injection (Fig. 6B). The fluorescence measured in the excised organs confirmed that a prevalent fraction of the HFn-CTX nanoconjugate not trapped in the tumor mass was rapidly sequestered by the liver, as previously described.^40,41^ Notably, the liver exhibited a detectable emission at all time points (Fig. 6C). Besides the liver, the lungs and the spleen showed the presence of the HFn-CTX nanoconjugate until 3 h post injection (Fig. 6C). Confocal imaging of tumor cryosections corroborated that the HFn-CTX nanoconjugate started to accumulate in the tumor mass early after injection and was no longer visible at 24 h (Fig. 7).

**Figure 6.**
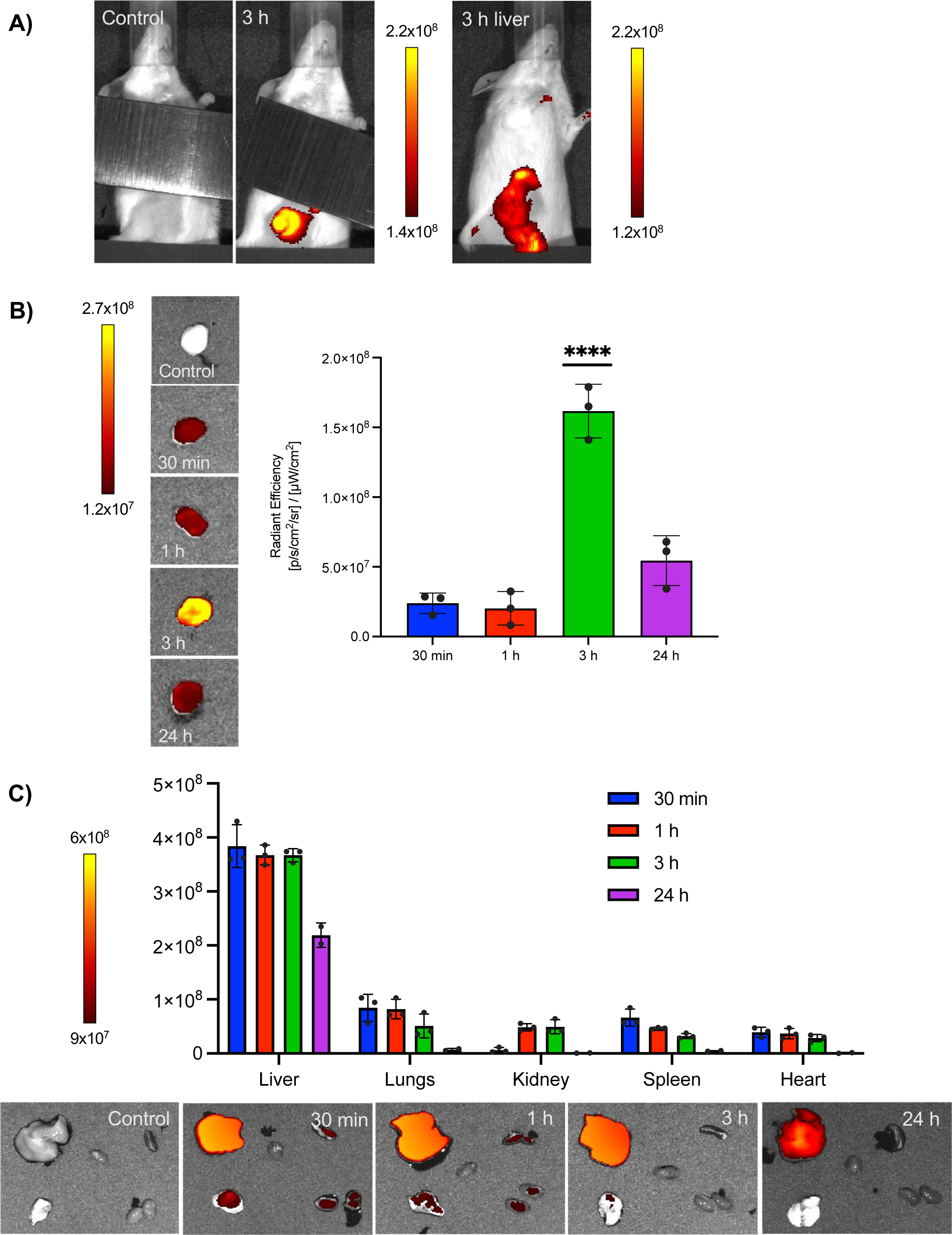
The HFn-CTX nanoconjugate displays efficient long-lasting tumor-targeting. **(A)** (*Left*) Representative epifluorescence (Epf) images of tumor-bearing mice taken 3 hours (3 h) after intravenous injection of 5 μg kg^-1^ HFn-AF647-CTX or buffer alone (Control). A ruler (visible as a gray bar in the pictures) was put on the belly to mask the signal deriving from the accumulation of the nanoconjugate within the liver. (*Right*) Representative Epf images of tumor-bearing mice taken 3 h after intravenous injection showing also the signal derived from the liver. **(B)** Epf of isolated MDA-MB-231 tumors and averaged Epf intensity of tumor ROI acquired 30 min, 1, 3, and 24 h after HFn-CTX injection. **(C)** Epf of isolated spleen (S), kidneys (K), liver (L), heart (H), lungs (Lu), and averaged Epf intensity of the ROI obtained after 30 min, 1, 3, and 24 h after HFn-CTX injection. The color scale in A, B, and C, indicates the averaged epifluorescence expressed as radiant efficiency [(p/sec/cm^2^/sr)/ (mW/cm^2^)], where p/sec/cm^2^/sr is the number of photons per second that leave a cm^2^ of tissue and radiate into a solid angle of one steradian (sr). Values reported in B and C are the mean ± SE of 3 independent samples for each experimental condition. **** = p < 0.0001 (One way ANOVA).

**Figure 7.**
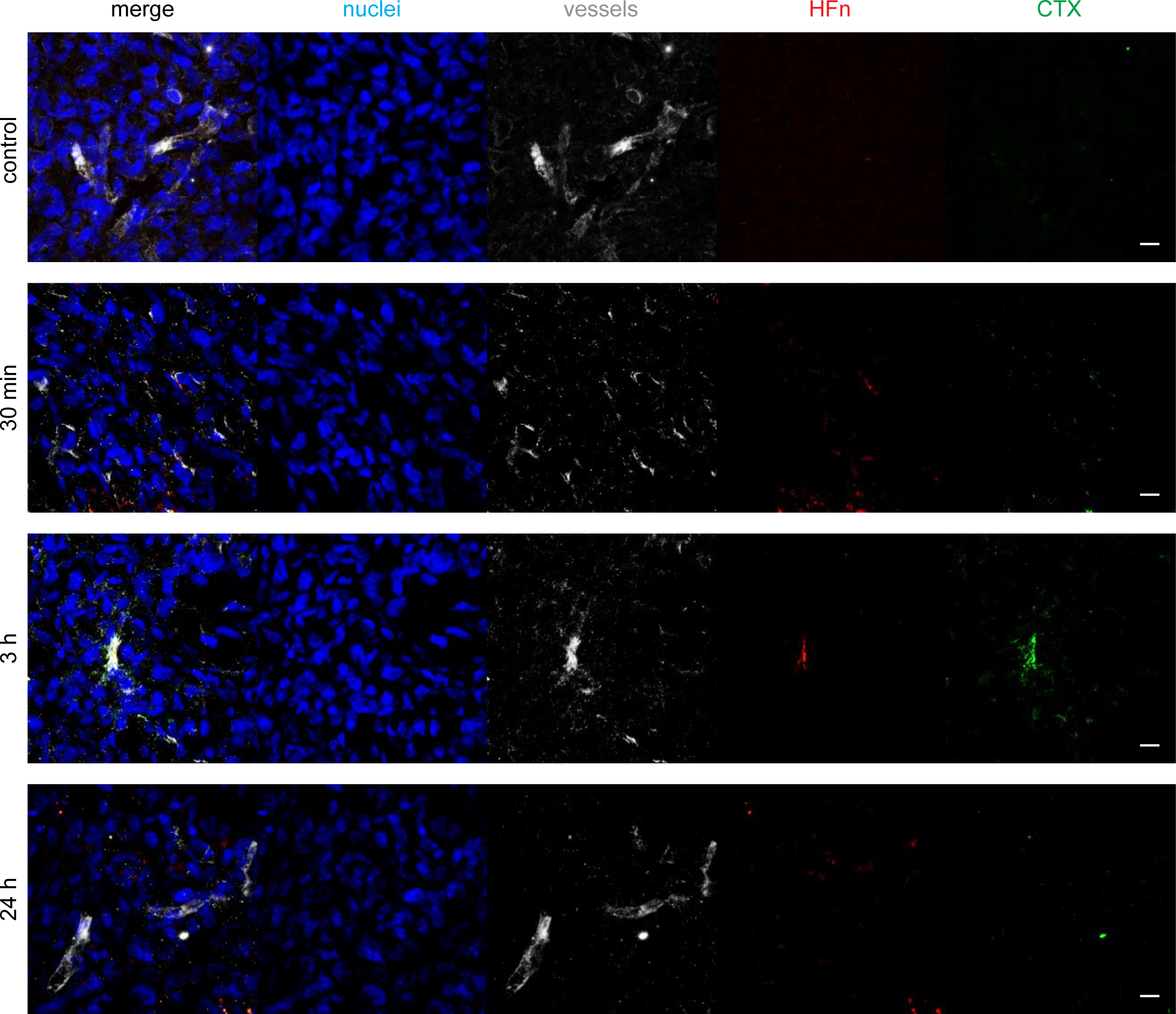
Intratumoral distribution of the HFn-CTX nanoconjugate. Representative images of 15 µm-thick cryosections of tumors excised 30 min, 1 h, 3 h, and 24 h after the injection of the nanoconjugate or PBS as a control (control). Confocal Z-stack images were acquired with a 0.5 µm step size. Stacks were summed together using ImageJ to obtain the images shown above. Background was subtracted using the rolling ball method with a radius of 4.0 pixels. Nuclei were stained with DAPI (blue), vessels with anti-CD31 antibody (gray), CTX conjugated on the surface of HFn with anti-human AF488 antibody (green). AF647-labelled HFn is shown in red. Scale bar, 10 µm.

### Bioavailability of the HFn-CTX nanoconjugate

Blood samples were collected from the retro-orbital plexus prior to injecting the HFn-CTX nanoconjugate and 30 min, 1 h, 3 h, and 24 h after, and the intensity of fluorescence was measured in these samples. The nanoconjugate signal progressively decreased over time (Fig. S5A), in accordance with its biodistribution and clearance. We also determined the concentration of the HFn-CTX nanoconjugate using a calibration curve (Fig. S5B) from which the percentage of injected dose (% of ID) in the plasma could be derived for each mouse. At 30 minutes post injection, the plasma retained only about 38% of the ID (Fig. 8A), showing that the HFn-CTX nanoconjugate readily reached both the organs and the tumor (Fig. S5C)

**Figure 8.**
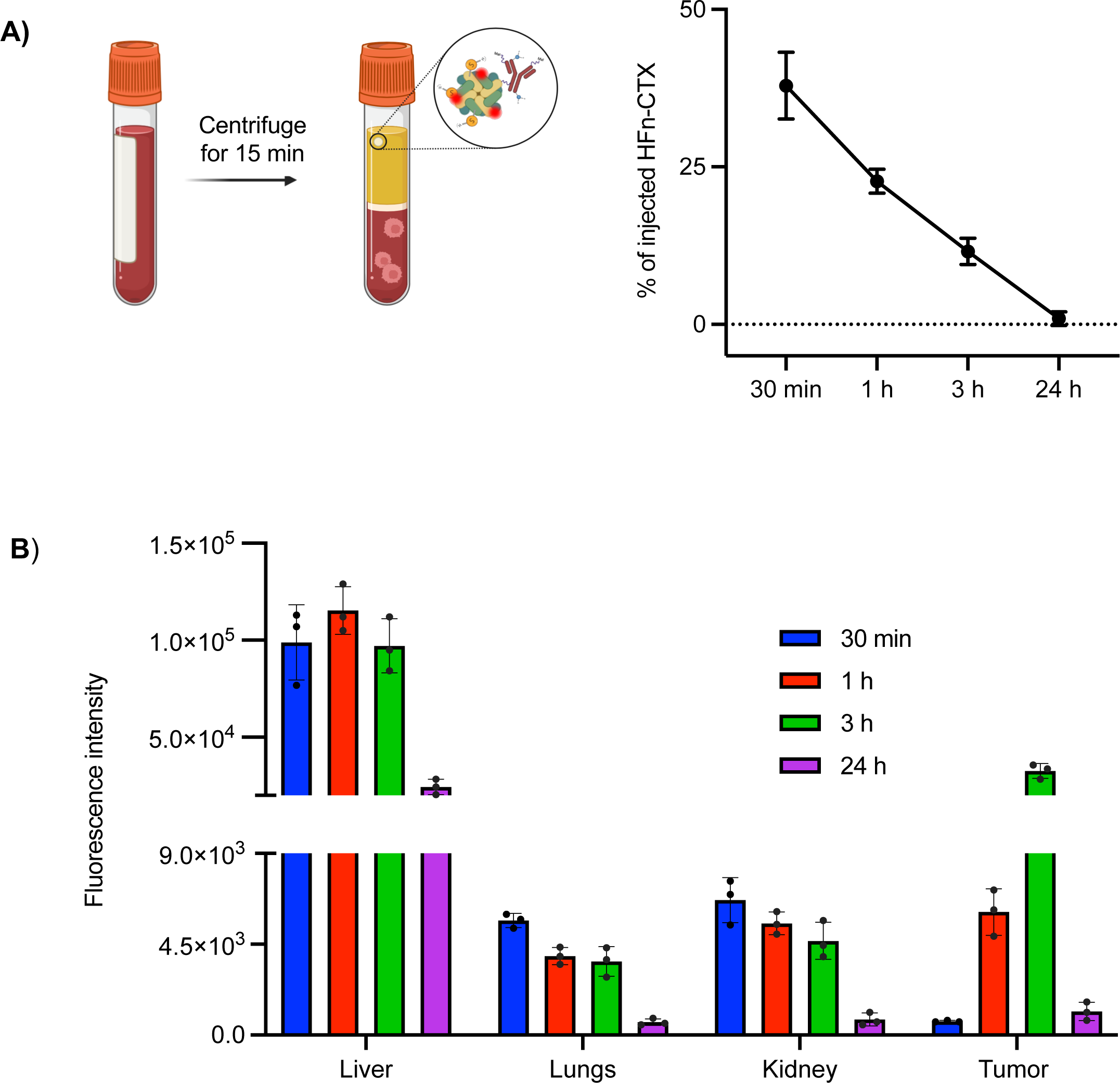
Distribution of the AF647-labeled HFn-CTX nanoconjugate in off-target organs and tumor homogenates. **(A)** Percentage of the injected dose (% of ID) of the AF647-labeled HFn-CTX nanoconjugate in mouse plasma. Reported values are mean ± SE of 3 different samples under each experimental condition. **(B)** Fluorescence intensity of the AF647-labeled HFn-CTX nanoconjugate measured in homogenated tumors and organs. Data represent mean ± SE of 3 different samples for each experimental condition.

To investigate the bioavailability of the nanoconjugate to the tumor and major organs, we exploited homogenized tissues (Fig. 8B). We found that the HFn-CTX nanoconjugates accumulated preferentially in the liver, as previously published for HFn alone and in agreement with the detoxification role of this organ. After 24 h, there was a consistent decrease in the fluorescence intensity in all organs tested (liver, kidneys, and lungs).

The tumor homogenates confirmed the accumulation of the HFn-CTX nanoconjugate peaked at 3 hours but revealed also a higher fluorescence signal already at 1 h post injection, which was probably underestimated in the *ex-vivo* analysis, due to its lower sensitivity. Confocal imaging of tumor cryosections corroborated this result: the HFn-CTX nanoconjugate was accumulated inside blood vessels at 1 and 3 h post injection (Fig. 7) and then presumably diffused across the tumor mass as it was no longer detectable at 24 h.

Importantly, the sum of these analyses showed that the tumor represents the second preferred site of accumulation and retention of the nanoconjugate, which thus appears to exhibit superior properties as compared to HFn nanoparticles.^42,43^

## Discussion

CTX is an anti-EGFR monoclonal antibody that has a proven potential in treating EGFR-positive cancers. However, primary or acquired mutations in the Ras-ERK and PI3K-AKT pathways confer resistance to CTX on most EGFR-positive tumors, such as glioblastoma and TNBC, for which limited molecular targets are available.

In this study, we developed a dual-targeting strategy that leverages HFn nanoconjugates to repurpose CTX for the treatment of CTX-resistant cancers. Our approach capitalizes on HFn to target the TfR1, which is highly expressed in nearly all solid tumors, and on the EGFR-binding ability and immune-mediated killing activity of CTX.

In this way, we enhanced the CTX’s anti-tumor efficacy on refractory cells: the ADCC response observed in TNBC spheroids treated with the HFn-CTX nanoconjugate was markedly stronger compared to CTX alone, suggesting that this dual-targeting approach effectively amplifies the immune-mediated anti-tumor response. Surprisingly, the nanoconjugate had no such effect on glioblastoma spheroids, suggesting the existence of unappreciated factors that allow the nanoconjugate to boost the CTX’s immune-mediated anti-tumor response.

We identified the cell-surface levels of EGFR and TfR1 as a key factor underlying this difference. The finding that the uptake of the nanoconjugate is faster in the U-87 MG cells than in MDA-MB-231 cells is in line both with our observations and evidence from the literature, as summarised below.

Firstly, U-87 MG cells show high levels of TfR1, which is known to undergo constitutive ligand-independent CME and recycling.^37^ Secondly, they have a relatively low amount of EGFR on the plasma membrane. This is relevant because, at odds with the TfR1, the endocytosis of EGFR is mainly driven by ligands that bind and activate the EGFR, thereby inducing its clathrin-dependent internalization. Thirdly, ligand-free, inactive EGFR is predominant in most tumor microenvironments, where the concentration of EGFR ligands is usually low.^44^ Fourthly, CTX outcompetes EGFR ligands and could stimulate the uptake of ligand-free EGFR via alternative secondary routes such as caveolin-mediated endocytosis.^45,46^ Fifthly, caveolin-mediated endocytosis is also kinetically slower than clathrin-mediated endocytosis, plays a negligible role in the uptake of nanoparticles, and might be negatively impacted by the nanoconjugate-mediated formation TfR1-EGFR clusters.^47,48^ This collective evidence likely explains why the HFn-CTX nanoconjugate lingers longer on the plasma membrane in MDA-MD-231 than in U-87 MG cells and thereby triggers ADCC only in the former. Anyway, the ability of HFn-CTX to overcome resistance to CTX in TNBC underscores the power of targeting multiple receptors to improve therapeutic effectiveness and specificity.

In addition to enhancing ADCC in TNBC spheroids, the HFn-CTX nanoconjugate exhibited favorable biophysical properties, including a hydrodynamic size of <30 nm, facilitating tumor penetration. Indeed, it showed superior biodistribution and efficient accumulation in TNBC tumors *in vivo*, possibly resulting from the combined effects of size and dual targeting. Of note, the modular nature of the HFn nanocage allows for potential functionalization with other therapeutic mAbs or biologics, broadening its applicability beyond TNBC.

In a broader perspective, our data strengthen the notion that nanotechnology may overcome the limitations of conventional systemic mAbs therapies, such as inadequate pharmacokinetics, poor tumor penetration, and off-target toxicity.

Although the therapeutic potential of the HFn-CTX nanoconjugate remains to be explored in full, we demonstrate the feasibility of repurposing CTX for TNBC therapy through a dual-targeting strategy that exploits HFn. Hence, the HFn-CTX nanoconjugate qualifies as a prime candidate for advanced nanomedicine applications to increase the clinical success of CTX-based therapies in TNBC and other refractory cancers.

## Materials and methods

### Reagents and antibodies

Mammalian cell culture media and reagents were from Euroclone. If not otherwise specified, all other chemicals were from Sigma-Aldrich.

Primary antibodies: CD44 (clone IM7; Biorad, MCA4703), CD31/PECAM-1 (Biotechne, AF3628), CD71 (clone D7G9X, Cell Signaling, 13113)^49,50^, EGFR (clone D38B1, Cell Signaling, 4267), β-Actin (clone D6A8, Cell Signaling, 8457), EGFR (clone 528, Merck, GR01), CD71 (clone MEM-75, Invitrogen, MA1-19137), Ferritin Heavy Chain (clone EPR3005Y, Abcam, ab75972).

Secondary antibodies: AF-conjugated secondary antibodies were from Thermo Fisher Scientific. HRP-conjugated secondary antibodies were from Cell Signalling (Goat anti-rabbit IgG, 1:2000, 7074; mouse anti-rabbit IgG conformation specific, 1:2000, 3678). Rabbit anti-Human IgG (H+L) HRP-linked Antibody (1:5000, Bethyl, A80-118P).

### Production and purification of HFn nanoparticles

Heavy chain apoferritin (HFn) was produced as previously described.^33^ Briefly, *Escherichia coli* ClearColi® BL21(DE3) strain (Lucigen Ltd. UK) carrying a pET30b plasmid encoding for HFn was grown at 37 °C in Luria Bertani medium supplemented with kanamycin until optical density (OD) at 600 nm reached 0.6. Then, the cells were induced with 1 mM of isopropyl β-D-1-thiogalactopyranoside (IPTG). After 3 hours, they were collected, washed with phosphate-buffered saline (PBS), and resuspended in a lysis buffer with lysozyme, DNase Benzonase and protease inhibitors. After the sonication, the crude extract was heated at 70 °C for 15 min and centrifuged. The supernatant was loaded onto diethylaminoethanol (DEAE) Sepharose anion exchange resin, pre-equilibrated with 2-(N-Morpholino)ethanesulfonic acid potassium salt (K-MES) 20 mM, pH 6.0. A stepwise NaCl gradient was used to elute the purified protein. The fractions were analyzed by Sodium Dodecyl Sulphate - PolyAcrylamide Gel Electrophoresis (SDS-PAGE) using 12% (v/v) polyacrylamide gels. The HFn-rich fractions were pulled, loaded on Amicon filters (100 kDa MWCO) and washed several times with PBS to remove traces of small protein contaminants and thereby obtain purer HFn. Protein concentration of the final preparation was determined using the Coomassie Plus Protein Assay Reagent (Thermo Fisher Scientific, MA, USA) and by measuring absorbance at 280 nm. Its purity was assessed by measuring the absorbance ratio 260/280 with a UV-vis spectrophotometer (NanoDrop™ 2000 Spectrophotometer, Thermo Fisher Scientific, MA, USA). HFn nanoparticles were stored at -80 °C in Tris-HCl 50 mM pH7.6, NaCl 150 mM, 10% glycerol to maximize their stability and solubility, as previously described.^51^

### Conjugation and purification of HFn-CTX nanoconjugates

The conjugation reaction was adapted from *Falvo et al*,.^32^ as briefly described below. Amine-containing CTX and sulfhydryl-containing HFn were covalently conjugated by means of a heterobifunctional crosslinker bearing N-hydroxysuccinimide (NHS) ester and maleimide (Mal) groups (Malhex-CONH-PEG-NHS, 5 kDa, Rapp Polymere Gmbh, Tubingen). CTX was reacted with a 40-fold molar excess of the crosslinker in phosphate buffer (PBS) pH 7.5 at room temperature for 1 h, and then unreacted species were first removed by washing with PBS buffer in 50 kDa centrifugal filter devices (Amicon, Millipore Corporate) and then on a size-exclusion chromatography (SEC) column (Zeba™ Spin Desalting Columns, Thermo Fisher Scientific, MA, USA). HFn molecules were added to PEGylated CTX at a final HFn:CTX molar ratio of 1:1 and incubated overnight at 4 °C under stirring. At the end of the reaction, large aggregates were removed by centrifugation (15,000xg for 15 min at 4 °C). The nanoconjugate-containing supernatant was collected and then purified through two SEC-FPLC steps, the former and the latter employing a Superdex 200 increase 10/300 GL column and a Superose 6 increase 10/300 GL column (GE Healthcare) equilibrated with PBS, respectively, and a flow rate of 0,2 mL min^−1^. The eluted fractions were then characterized with SDS-PAGE, western blot, and dynamic light scattering (DLS) analyses. The fractions containing the HFn-CTX nanoconjugates were concentrated on Amicon filters (100 kDa MWCO), if deemed necessary. HFn-CTX nanoconjugates were kept at 4 °C in PBS after assessing their stability by means of UV-VIS spectrometry and/or centrifugation.

### Labeling of HFn-CTX and HFn nanoparticles

Double-labeled nanoconjugates were generated using Alexa Fluor™-750 NHS Ester (AF750, Thermo Fisher Scientific, MA, USA) and Alexa Fluor™-647-NHS-ester (AF647, Thermo Fisher Scientific, MA, USA).

To label HFn with AF647, a Zeba™ Spin desalting column (7 kDa) was used to exchange the HFn storage buffer for 0.1 M NaHCO_3_ pH 8.0, as per manufacturer’s instructions. AF-647-NHS-ester (20 mg mL^-1^ in DMSO) (Thermo Fisher Scientific, Waltham, MA, USA) was then added to the HFn solution (5 mg mL^-1^) to obtain a molar dye/protein ratio of 50. This mix was stirred at RT for 30 min and then at 4 °C overnight. The unreacted dye was removed with a Zeba™ Dye and Biotin Removal Columns (7 kDa, Thermo Fisher Scientific, Waltham, MA, USA) using 50 mM potassium phosphate, pH 7.2, and 150 mM NaCl as the final buffer. The degree of labeling was calculated as the molar ratio between dye and protein, based on UV–Vis spectroscopy. To this end, 280 nm and 647 nm absorbances were used for the quantification of the protein and the dye, respectively, subtracting the contribution of the dye at 280 nm using the following formula:

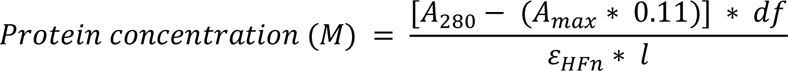

where Amax is absorbance (A) of a dye solution measured at the wavelength maximum (λmax) for the dye molecule, 0.03 is the correction factor recommended by the supplier to remove the contribution of Alexa Fluor™-647-NHS-ester to the 280 nm absorbance, df is the dilution factor, which represent the extent (if any) to which the protein:dye sample was diluted for the absorbance measurement, l (optical path length) is 0.1 cm, εHFn is 458,340 M^-1^ cm^-1^ (calculated with ProtParam)^52^.

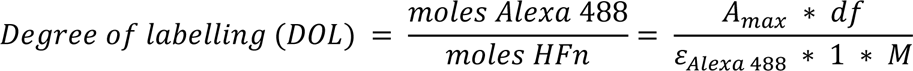

where εAlexa647 (molar extinction coefficient of Alexa Fluor™-647-NHS-ester) is 239,000 M^-1^ cm^-1^, and M is the protein concentration. HFn-647 was stored at -80 °C in a previously described buffer.^53^

CTX was labeled with AF750 using the same procedure and similar reaction conditions. In this case, AF-750-NHS-ester (10 mg mL^-1^ in DMSO) (Thermo Fisher Scientific, Waltham, MA, USA) was added to the CTX solution (5 mg mL^-1^) to obtain a molar dye/protein ratio of 10. Before starting the reaction, a 40-fold molar excess of the PEG crosslinker was added to the reaction mix because both AF750 and PEG crosslinker compete to bind to the amine groups of CTX. Then the reaction was kept under stirring for 2.5 hours at RT. At the end of the incubation, the unreacted dye was removed as described above. Labeled CTX was further purified using SEC-FPLC as described above. Quantification of the protein and the dye was performed using the formula reported above (0.04 is the correction factor recommended by the supplier to remove the contribution of the AF-750-NHS-ester from the 280 nm absorbance, εAlexa750 is 240,000 and εCTX is 217,440 M^-1^ cm^-1^ as calculated by ProtParam).

### Cell culture

U-87 MG (ATCC, HTB-14) was cultured in Dulbecco’s Modified Eagle’s Medium (DMEM) supplemented with 10% fetal bovine serum (FBS), 2mM L-glutamine, penicillin (50 UI mL^-1^) and streptomycin (50 mg mL^-1^). MDA-MB-231 cells (Amsbio, SC059-Puro) were cultured in Minimum Essential Medium (MEM) supplemented with 10% FBS, 2 mM L-glutamine, penicillin (50 UI mL^−1^) and streptomycin (50 mg mL^−1^). All cell lines were maintained at 37 °C in a humidified atmosphere containing 5% CO_2_ and passaged prior to confluence. Cells were regularly tested for mycoplasma, and all experiments were performed on mycoplasma-free cells.

### Spheroids generation

U-87 MG cells were detached with trypsin-EDTA and cell number was determined using a Bürker chamber. The cell suspension was diluted to 6x10^4^ cells mL^-1^ in complete medium. A volume of 100 μl of the cell suspension (6000 cells) was added into each well of an ultra-low attachment Nunclon™ Sphera™ 96-well plate (Thermo Fisher Scientific, MA, USA). Spheroid formation was initiated by spinning the plates at 340xg for 10 min using an Eppendorf 5810 centrifuge (Eppendorf AG, Hamburg) with swinging buckets. The plates were incubated at 37 °C and 5% CO_2_ in a humidified incubator for 5 days.

MDA-MB-231 cells were detached and counted as above. The cell suspension was diluted to 2.5x10^4^ cells mL^-1^ in ice-cold medium. Cultrex® (3-D Culture Matrix Reduced Growth Factor BME, R&D SYSTEM) was thawed on ice overnight and added with ice-cold pipette tips at a final concentration of 2.5% to the cell suspension. A volume of 200 μl of the cell suspension (5000 cells) was added into each well of an ultra-low attachment Nunclon™ Sphera™ 96-well plate (Thermo Fisher Scientific, MA, USA). Spheroid formation was initiated as above, and the plates were incubated at 37 °C and 5% CO_2_ in a humidified incubator for 5 days. Spheroid area was measured at the Incucyte (Sartorius) using the built-in spheroid analysis module.

### Isolation of effector cells

Peripheral blood mononuclear cells (PBMCs) were obtained by centrifugation on Ficoll-Paque of blood samples (50 mL) from healthy human donors (408xg without brakes for 30 min. at RT). The PBMCs layer was carefully transferred into a new 50 mL tube, diluted with PBS, and centrifuged at 1,400 rpm for 6 min at RT. Then, the supernatant was discarded to remove platelets, and this procedure was repeated four times decreasing the centrifugation speed down to 1,000 rpm. Washed PBMCs were resuspended in RPMI-1640 medium supplemented with 10% of heat-inactivated FBS and 1,000 U mL^-1^ IL-2 (BioLegend, San Diego, CA, USA) for 24 h at 37 °C. This procedure was approved by the Ethical Committee of the University of Milano-Bicocca after the submission of the project together with informed consent by the healthy volunteers (prot. 0138485/21, November 15^th^, 2021).

### Caspase activation assay

For endpoint assays, apoptosis of target cells was evaluated using the Caspase-Glo® 3/7 3D (Promega Corporation, Madison, WI, USA). Spheroids were first washed and transferred in FBS-free RPMI culture medium, prior to treatment with free CTX, or HFn-CTX nanoconjugates (10 μg mL^-1^), or HFn (34 μg mL^-1^), all diluted in FBS-free RPMI culture medium to reach a final volume of 0.2 mL. Note that HFn, was given at 34 μg mL^−1^ because 10 μg mL^−1^ of CTX in the nanoconjugate formulation correspond to 34 μg mL^−1^ of HFn, because the two protein species have a molar ratio of 1:1. After incubation at 4 °C for 30 min, IL-2-activated PBMCs were added onto the target cells at an effector:target (E:T) ratio of 2:1 and incubated at 37 °C for 24 or 48 h. Then, 100 μL of Caspase-Glo® 3/7 3D Reagent were added to each well, and the plate was incubated at RT for at least 30 minutes. Luminescence, which is proportional to the amount of caspase activity, was measured with an EnSight™ multimode plate reader (Perkin Elmer, Walthman, MA, USA).

### HFn-CTX and HFn uptake assays

U-87 MG and MDA-MB-231 cells were seeded onto 4-compartment 35 mm glass bottom Petri dishes (Greiner Bio-One) at a density of 130,000 cells and 60,000 cells per compartment, respectively, in 0.5 mL of complete medium. On the following day, the medium was replaced with 0.2 mL of fresh complete medium containing HFn-AF647-CTX-AF750 (double-labelled nanoparticles) or HFn-AF647 (single-labelled nanoparticles). The cells were then incubated for 0, 2, 15, 30, 60, and 120 min at 37°C.

For the 2D cultures, the growth medium was removed, and the cells were washed twice with pre-warmed PBS. Subsequently, they were fixed and immunostained as previously described.^54,55^ Nuclei were stained with Hoechst (5 µg mL^−1^) for 5 minutes at RT. Cells were then rinsed once with PBS before being imaged using a fluorescence microscope (Thunder Imager Live Cells, LEICA Microsystem).

For the spheroid cryosections, a Dakopen was used to draw a circle around the sectioned spheroids to create a water-repellent barrier. Glass slides were washed in PBS prior to adding the blocking and permeabilization buffer (0,2% BSA + 0,1% Triton-X100 in PBS) at RT for 10 min. After three washes with PBS, immunostaining was performed by incubating slices with primary antibodies diluted in PBS, 1% BSA at RT for 1 hour. Next, spheroid slices were rinsed thrice with PBS prior to the addition of secondary antibodies diluted in PBS, 1% BSA at RT for 45 minutes in the dark. After three washes with PBS, nuclei were stained with Hoechst (5 µg mL^−1^) by adding 50 µL on each drawn circle and incubating at RT for 5 minutes. Spheroid sections were then washed thrice with PBS and covered with a coverslip on which a drop of FluorSave Reagent (Millipore Corporation) was deposited. Mounted cryosections were dried at 37 °C for 20 minutes and then imaged using a fluorescence microscope (Thunder Imager Live Cells, LEICA Microsystem). Identical settings were used to acquire different experimental groups.

### Embedding and cryosectioning of spheroids

Spheroids were fixed in pre-warmed 4% paraformaldehyde in PEM buffer (80 mM PIPES pH 6.8, 5 mM EGTA, 2 mM MgCl_2_)^51,55^ for 2 h at RT (200 µL in each 96 wells), washed once with PBS w/o Ca^2+^ and Mg^2+^, and then dehydrated with 30% sucrose for 2 h at RT. At the end of the incubation, the spheroids were cryopreserved in OCT compound (R0030, Histo-Line laboratories) exploiting a freezing bath filled with ethanol and dry-ice. Frozen samples were then stored at -80 °C until sectioning. Before sectioning, embedded spheroids were left at -20 °C for at least 1 h. Slices of 7 µm were cut using a MC4000 cryostat (Histo-Line laboratories) and collected on SuperFrost Plus glass slides (HL26766, Histo-Line laboratories), which were stored at -80 °C for further use.

### Cell-surface EGFR and TfR1 expression levels

U-87 MG and MDA-MB-231 cells were co-seeded onto 35 mm glass-bottom Petri dishes (Greiner Bio-One), each at a density of 70,000 cells, in 1.25 mL of complete medium. Before plating, MDA-MB-231 cells were stained with Vybrant^TM^ CFDA SE Cell Tracer Kit (Thermo Fisher Scientific, Waltham, MA, USA) according to the manufacturer’s instructions. On the following day, co-cultures were fixed and immunostained using either anti-EGFR or anti-TfR1 antibodies. To stain for total or only cell-surface receptors, cells were blocked with or without Triton as a permeabilizing agent, respectively. Nuclei were stained with Hoechst (5 µg mL^−1^) for 5 minutes at RT. Non-permeabilized samples stained for a cytosolic marker, typically actin, provided a control for the integrity of the specimens and and for the specific detection of cell-surface signals (not shown). Cells were then rinsed once with PBS prior to imaging at a Thunder Imager Live Cells microscope (LEICA Microsystem), using identical acquisition settings. The images were then analyzed with Cellprofiler to quantify cell-surface EGFR and TfR1 levels.

### Image processing and analysis

Colocalization analysis was performed using the Mander’s colocalization plugin of ImageJ (National Institutes of Health, Bethesda, MD, USA). This coefficient measures the pixel-by-pixel covariance in the signal levels of two images.^56^ The background was subtracted before the analysis using a rolling ball of 15.0 pixels. The colocalization analysis was performed on three sections for which Manders’ coefficients were calculated with automatic threshold, which is e algorithmically determined by optimizing the separation between signal and noise. This approach reduces the influence of artifacts and nonspecific signal intensities, providing a more realistic and robust representation of molecular colocalization (tM = 0 means no co-localization, tM = 1 means completed co-localization). For display, images of the cryosections were contrasted in Adobe photoshop, using identical settings.

Cellprofiler was used to write bespoke pipelines for the automated analysis of spheroids’ cryosections and cell-surface receptor levels.^57^

### Western blot

To quantify TfR1 and EGFR expression, MDA-MB-231 and U-87 MG cells were washed three times with ice-cold PBS, detached from Petri dishes with Trypsin (0,05%), and collected by centrifugation. They were then lysed with RIPA buffer supplemented with Halt™ Protease Inhibitor Cocktail (ThermoFisher). The lysates were incubated at 4 °C for 30 minutes on a rocking wheel and then centrifuged at 13000xg at 4 °C for 15 minutes. Proteins in the cleared lysates were quantified using the bicinchoninic acid assay (BCA). Fifteen µg of total proteins were separated by SDS/PAGE, and then transferred onto a PVDF membrane (0.45 µm pore size; Whatman, GE Healthcare Europe) as previously described.^53,58^ Membranes were blocked in TBS supplemented with Tween 20 0,1% and BSA 5% for 1h at RT and incubated with primary antibodies according to the manufacturer’s instructions, followed by HRP-conjugated secondary antibodies. Signal was acquired at ChemiDoc Imaging System (Biorad, USA).

### Densitometry

To estimate band intensities, non-saturated exposures of western blots were subjected to densitometric analyses using ImageJ. Normalized TfR1 and EGFR expression was determined as follows: the intensities of TfR1 or EGFR, and those of a housekeeping protein (e.g., actin) were determined by densitometry, and the TfR1 or EGFR/actin ratio was calculated as previously described.^51^

### Tumour xenografts

Female NOD/scid mice purchased from Envigo (Bresso, Italy) were housed in a fully equipped facility according to the guidelines of the Italian Ministry of Health. Experiments were conducted under an approved protocol (authorization n° 92/2022-PR). Luciferase-expressing MDA-MB-231 cells were cultured in DMEM high glucose supplemented with 10% fetal bovine serum, 2 mM L-glutamine, penicillin (50 IU mL^−1^) and streptomycin (50 mg mL^−1^), 0.1 mM MEM Non-essential Amino acids (NEAA). A 1:1 mixture comprising 2x10^6^ cells and Matrigel (Corning® Matrigel® Growth Factor Reduced (GFR) Basement Membrane Matrix, Phenol Red-free, LDEV-free) was inoculated subcutaneously into the mouse mammary fat pad in a final volume of 100 μL. The health and behavior of these mice were observed daily, and the weight was monitored every three days until the tumors reached a volume around 100 mm^3^ (typically after 3 weeks of inoculation) as measured with a caliper.

Tumours’ dimension was also evaluated by means of the bioluminescence (BLI) signal *in vivo*, which was measured using an IVIS Lumina II imaging system (Perkin Elmer), with an exposure time of one minute. Twenty-four hours before injecting the nanoparticles, BLI images were acquired 5, 8, and 10 min after peritoneal injection of 260 μg kg^-1^ luciferin (D-Luciferin potassium salt, Perkin Elmer). Bioluminescence measurements were used to divide mice into experimental groups with a similar tumor burden.

### Tumour targeting and biodistribution of HFn-CTX *in vivo* and *ex vivo*

To study the biodistribution of HFn-CTX in mice, the nanoconjugate was labeled with AF647. NOD/SCID mice bearing luciferase-expressing MDA-MB-231 cell-derived tumours were immobilized in a restrainer (2 biological instruments). Next, HFn-AF647-CTX (5 mg kg^-1^ body weight) was injected into the tail vein with a 26G insulin needle, with PBS being used as a control. Epifluorescence images of anesthetized mice were obtained 30 minutes, 1h, 3h, 24h post-injection at the IVIS system (Perkin Elmer), keeping the animals at 37 °C. Images were acquired with a Cy5.5 emission filter, while excitation was scanned from 570 to 640, and autofluorescence was removed by spectral unmixing. The instrument setup for FLI acquisition is Fstop 2 and medium binning, T exposure 30 sec. After being imaged, those mice were sacrificed, and organs were dissected and imaged with the IVIS system at RT. All epifluorescence (Epf) intensity values were normalized after subtracting Epf values obtained from mice injected with PBS, and results were expressed as mean ± s.e.

Among off-target organs, liver, lungs, and kidneys exhibited the higher accumulation of the nanoconjugate as determined by IVIS analysis. Tissue samples at a concentration of 250 mg mL^-1^ in PBS were mechanically homogenized using a TissueLyser II (QIAGEN) at a frequency of 30 s^-1^ and beads for 15 minutes at 4 °C. Fluorescence at 665 nm was measured using an EnSight™ multimode plate reader (Perkin Elmer, Waltham, MA, USA) and from 100 µL of each suspension. This signal, upon subtraction of the blank (i.e., corresponding control mouse tissue), was correlated with the concentration of nanoconjugate by spiking the corresponding control mouse tissues with known concentrations of HFn-AF647-CTX. By measuring the weight of each organ and knowing the injected dose of nanoconjugate, we could calculate the percentage of injected nanoconjugate present at different time points (ID%, Injected Dose Percentage).

Tumors were processed using the same method at a concentration of 200 mg mL^-1^ using RIPA buffer for lysis.

Plasma from each mouse was obtained by spinning at 2,980xg for 15 minutes at 4 °C blood samples collected prior to sacrifice in heparinized tubes. Fluorescence at 665 nm was measured and correlated with the concentration of nanoconjugate as described above.

### Statistical analysis

Statistical analyses were conducted using adequate tests, including T test, one-way Anova, and two-way Anova. Graphs show mean values ± standard deviation (s.d.) or standard errors (s.e.). All tests assumed normal distribution and the statistical significance threshold was set at p<0.05.

## Author contributions

D.P. and M.I designed and supervised the project. L.B., A.B., L.S., S.G. and C.B. performed the *in vitro* experiments. L.B., L.S., A.B. and L.F. performed the *in vivo* studies. G.F., L.B. and A.B. produced the recombinant HFn nanoparticles. L.B., A.B.,

M.G. and L.M. synthesized and characterized HFn-CTX nanoconjugates. L.B., L.S., A.B., L.F., S.G. and M.I analyzed the results. M.I. and L.B. wrote the first draft of the manuscript. M.I., L.B., L.S., D.P., and M.C. revised the manuscript. All authors read and approved the final version.

## Supporting information

This manuscript is accompanied by four supplementary tables (Table S1-4) and 5 supplementary figures (Figure S1-5):

- Table S1. Physicochemical properties of HFn-CTX nanoconjugates.

- Table S2. Optimization of HFn labeling with AF488.

- Table S3. Optimization of CTX labeling with AF647.

- Table S4. Optimization of HFn labeling with AF647.

- Table S5. Optimization of CTX labeling with AF750.

- Figure S1. Two-step purification of the HFn-CTX nanoconjugate by SEC-FPLC.

- Figure S2. Distribution of the HFn-CTX nanoconjugate and HFn nanoparticles within GBM and TNBC spheroids.

- Figure S3. Large-field images of the uptake of the HFn-CTX nanoconjugate or HFn nanoparticles in U-87 MG and MDA-MB-231 cells.

- Figure S4. Large-field images of cell-surface EGFR and TfR1 levels in U-87 MG and MDA-MB-231 co-cultures.

- Figure S5. Biodistribution of the HFn-CTX nanoconjugate in mice.

## Acknowledgments

The research leading to these results has received funding from AIRC under IG 2018 – ID. 21565 project – P.I. D. Prosperi. Support to this work was also provided by the European Union – NextGenerationEU through the Italian Ministry of University and Research under PNRR – M4C2-I1.3 Project PE_00000019 “HEAL ITALIA” to D. Prosperi CUP H43C22000830006 of University of Milano-Bicocca.

## Notes

### Competing Interest Statement

The authors have declared no competing interest.

## References

(1) Harbeck, N.; Penault-Llorca, F.; Cortes, J.; Gnant, M.; Houssami, N.; Poortmans, P.; Ruddy, K.; Tsang, J.; Cardoso, F. Breast Cancer. Nat Rev Dis Primers 2019, 5 (1), 66. 10.1038/s41572-019-0111-2.

(2) Noone, A.-M.; Cronin, K. A.; Altekruse, S. F.; Howlader, N.; Lewis, D. R.; Petkov, V. I.; Penberthy, L. Cancer Incidence and Survival Trends by Subtype Using Data from the Surveillance Epidemiology and End Results Program, 1992–2013. Cancer Epidemiology, Biomarkers & Prevention 2017, 26 (4), 632–641. 10.1158/1055-9965.EPI-16-0520.

(3) Singh, D. D.; Yadav, D. K. TNBC: Potential Targeting of Multiple Receptors for a Therapeutic Breakthrough, Nanomedicine, and Immunotherapy. Biomedicines 2021, 9 (8), 876. 10.3390/biomedicines9080876.

(4) Lee, K.-L.; Kuo, Y.-C.; Ho, Y.-S.; Huang, Y.-H. Triple-Negative Breast Cancer: Current Understanding and Future Therapeutic Breakthrough Targeting Cancer Stemness. Cancers (Basel) 2019, 11 (9), 1334. 10.3390/cancers11091334.

(5) Won, K.-A.; Spruck, C. Triple-negative Breast Cancer Therapy: Current and Future Perspectives (Review). Int J Oncol 2020, 57 (6), 1245–1261. 10.3892/ijo.2020.5135.

(6) Nedeljković, M.; Damjanović, A. Mechanisms of Chemotherapy Resistance in Triple-Negative Breast Cancer-How We Can Rise to the Challenge. Cells 2019, 8 (9), 957. 10.3390/cells8090957.

(7) Masuda, H.; Zhang, D.; Bartholomeusz, C.; Doihara, H.; Hortobagyi, G. N.; Ueno, N. T. Role of Epidermal Growth Factor Receptor in Breast Cancer. Breast Cancer Res Treat 2012, 136 (2), 10.1007/s10549-012-2289–9. 10.1007/s10549-012-2289-9.

(8) Corkery, B.; Crown, J.; Clynes, M.; O’Donovan, N. Epidermal Growth Factor Receptor as a Potential Therapeutic Target in Triple-Negative Breast Cancer. Annals of Oncology 2009, 20 (5), 862–867. 10.1093/annonc/mdn710.

(9) Blick, S. K. A.; Scott, L. J. Cetuximab: A Review of Its Use in Squamous Cell Carcinoma of the Head and Neck and Metastatic Colorectal Cancer. Drugs 2007, 67 (17), 2585–2607. 10.2165/00003495-200767170-00008.

(10) Baysal, H.; De Pauw, I.; Zaryouh, H.; Peeters, M.; Vermorken, J. B.; Lardon, F.; De Waele, J.; Wouters, A. The Right Partner in Crime: Unlocking the Potential of the Anti-EGFR Antibody Cetuximab via Combination With Natural Killer Cell Chartering Immunotherapeutic Strategies. Front. Immunol. 2021, 12. 10.3389/fimmu.2021.737311.

(11) Lo Nigro, C.; Ricci, V.; Vivenza, D.; Monteverde, M.; Strola, G.; Lucio, F.; Tonissi, F.; Miraglio, E.; Granetto, C.; Fortunato, M.; Merlano, M. C. Evaluation of Antibody-Dependent Cell-Mediated Cytotoxicity Activity and Cetuximab Response in KRAS Wild-Type Metastatic Colorectal Cancer Patients. World J Gastrointest Oncol 2016, 8 (2), 222–230. 10.4251/wjgo.v8.i2.222.

(12) Cai, W.-Q.; Zeng, L.-S.; Wang, L.-F.; Wang, Y.-Y.; Cheng, J.-T.; Zhang, Y.; Han, Z.-W.; Zhou, Y.; Huang, S.-L.; Wang, X.-W.; Peng, X.-C.; Xiang, Y.; Ma, Z.; Cui, S.-Z.; Xin, H.-W. The Latest Battles Between EGFR Monoclonal Antibodies and Resistant Tumor Cells. Front Oncol 2020, 10, 1249. 10.3389/fonc.2020.01249.

(13) Rodríguez-Nava, C.; Ortuño-Pineda, C.; Illades-Aguiar, B.; Flores-Alfaro, E.; Leyva-Vázquez, M. A.; Parra-Rojas, I.; del Moral-Hernández, O.; Vences-Velázquez, A.; Cortés-Sarabia, K.; Alarcón-Romero, L. del C. Mechanisms of Action and Limitations of Monoclonal Antibodies and Single Chain Fragment Variable (scFv) in the Treatment of Cancer. Biomedicines 2023, 11 (6), 1610. 10.3390/biomedicines11061610.

(14) Cruz, E.; Kayser, V. Monoclonal Antibody Therapy of Solid Tumors: Clinical Limitations and Novel Strategies to Enhance Treatment Efficacy. Biologics 2019, 13, 33–51. 10.2147/BTT.S166310.

(15) Nguyen, P. V.; Hervé-Aubert, K.; David, S.; Lautram, N.; Passirani, C.; Chourpa, I.; Aubrey, N.; Allard-Vannier, E. Targeted Nanomedicine with Anti-EGFR scFv for siRNA Delivery into Triple Negative Breast Cancer Cells. European Journal of Pharmaceutics and Biopharmaceutics 2020, 157, 74–84. 10.1016/j.ejpb.2020.10.004.

(16) Venugopal, V.; Krishnan, S.; Palanimuthu, V. R.; Sankarankutty, S.; Kalaimani, J. K.; Karupiah, S.; Kit, N. S.; Hock, T. T. Anti-EGFR Anchored Paclitaxel Loaded PLGA Nanoparticles for the Treatment of Triple Negative Breast Cancer. In-Vitro and in-Vivo Anticancer Activities. PLOS ONE 2018, 13 (11), e0206109. 10.1371/journal.pone.0206109.

(17) Liao, W.-S.; Ho, Y.; Lin, Y.-W.; Naveen Raj, E.; Liu, K.-K.; Chen, C.; Zhou, X.-Z.; Lu, K.-P.; Chao, J.-I. Targeting EGFR of Triple-Negative Breast Cancer Enhances the Therapeutic Efficacy of Paclitaxel- and Cetuximab-Conjugated Nanodiamond Nanocomposite. Acta Biomaterialia 2019, 86, 395–405. 10.1016/j.actbio.2019.01.025.

(18) Lammers, T. Nanomedicine Tumor Targeting. Advanced Materials 2024, 36 (26), 2312169. 10.1002/adma.202312169.

(19) Mazzucchelli, S.; Truffi, M.; Baccarini, F.; Beretta, M.; Sorrentino, L.; Bellini, M.; Rizzuto, M. A.; Ottria, R.; Ravelli, A.; Ciuffreda, P.; Prosperi, D.; Corsi, F. H-Ferritin-Nanocaged Olaparib: A Promising Choice for Both BRCA-Mutated and Sporadic Triple Negative Breast Cancer. Sci Rep 2017, 7 (1), 7505. 10.1038/s41598-017-07617-7.

(20) Shen, Y.; Li, X.; Dong, D.; Zhang, B.; Xue, Y.; Shang, P. Transferrin Receptor 1 in Cancer: A New Sight for Cancer Therapy. Am J Cancer Res 2018, 8 (6), 916– 931.

(21) Fontana, F.; Esser, A. K.; Egbulefu, C.; Karmakar, P.; Su, X.; Allen, J. S.; Xu, Y.; Davis, J. L.; Gabay, A.; Xiang, J.; Kwakwa, K. A.; Manion, B.; Bakewell, S.; Li, S.; Park, H.; Lanza, G. M.; Achilefu, S.; Weilbaecher, K. N. Transferrin Receptor in Primary and Metastatic Breast Cancer: Evaluation of Expression and Experimental Modulation to Improve Molecular Targeting. PLoS One 2023, 18 (12), e0293700. 10.1371/journal.pone.0293700.

(22) Singh, M.; Mugler, K.; Hailoo, D. W.; Burke, S.; Nemesure, B.; Torkko, K.; Shroyer, K. R. Differential Expression of Transferrin Receptor (TfR) in a Spectrum of Normal to Malignant Breast Tissues: Implications for in Situ and Invasive Carcinoma. Appl Immunohistochem Mol Morphol 2011, 19 (5), 417–423. 10.1097/PAI.0b013e318209716e.

(23) He, J.; Fan, K.; Yan, X. Ferritin Drug Carrier (FDC) for Tumor Targeting Therapy. J Control Release 2019, 311–312, 288–300. 10.1016/j.jconrel.2019.09.002.

(24) Rizzuto, M. A.; Magro, R. D.; Barbieri, L.; Pandolfi, L.; Sguazzini-Viscontini, A.; Truffi, M.; Salvioni, L.; Corsi, F.; Colombo, M.; Re, F.; Prosperi, D. H-Ferritin Nanoparticle-Mediated Delivery of Antibodies across a BBB in Vitro Model for Treatment of Brain Malignancies. Biomater. Sci. 2021, 9 (6), 2032–2042. 10.1039/D0BM01726D.

(25) Fan, K.; Jia, X.; Zhou, M.; Wang, K.; Conde, J.; He, J.; Tian, J.; Yan, X. Ferritin Nanocarrier Traverses the Blood Brain Barrier and Kills Glioma. ACS Nano 2018, 12 (5), 4105–4115. 10.1021/acsnano.7b06969.

(26) Huang, C.-W.; Chuang, C.-P.; Chen, Y.-J.; Wang, H.-Y.; Lin, J.-J.; Huang, C.-Y.; Wei, K.-C.; Huang, F.-T. Integrin Α2β1-Targeting Ferritin Nanocarrier Traverses the Blood–Brain Barrier for Effective Glioma Chemotherapy. Journal of Nanobiotechnology 2021, 19 (1), 180. 10.1186/s12951-021-00925-1.

(27) Deng, G.; Li, Y.; Liang, N.; Hu, P.; Zhang, Y.; Qiao, L.; Zhang, Y.; Xie, J.; Luo, H.; Wang, F.; Chen, F.; Liu, F.; Xu, D.; Zhang, J. Ferritin in Cancer Therapy: A Pleiotropic Tumoraffin Nanocage-Based Transport. Cancer Medicine 2023, 12 (10), 11570–11588. 10.1002/cam4.5778.

(28) Khoshnejad, M.; Greineder, C. F.; Pulsipher, K. W.; Villa, C. H.; Altun, B.; Pan, D. C.; Tsourkas, A.; Dmochowski, I. J.; Muzykantov, V. R. Ferritin Nanocages with Biologically Orthogonal Conjugation for Vascular Targeting and Imaging. Bioconjug Chem 2018, 29 (4), 1209–1218. 10.1021/acs.bioconjchem.8b00004.

(29) Truffi, M.; Fiandra, L.; Sorrentino, L.; Monieri, M.; Corsi, F.; Mazzucchelli, S. Ferritin Nanocages: A Biological Platform for Drug Delivery, Imaging and Theranostics in Cancer. Pharmacol Res 2016, 107, 57–65. 10.1016/j.phrs.2016.03.002.

(30) Clark, M. J.; Homer, N.; O’Connor, B. D.; Chen, Z.; Eskin, A.; Lee, H.; Merriman, B.; Nelson, S. F. U87MG Decoded: The Genomic Sequence of a Cytogenetically Aberrant Human Cancer Cell Line. PLOS Genetics 2010, 6 (1), e1000832. 10.1371/journal.pgen.1000832.

(31) Hollestelle, A.; Elstrodt, F.; Nagel, J. H. A.; Kallemeijn, W. W.; Schutte, M. Phosphatidylinositol-3-OH Kinase or RAS Pathway Mutations in Human Breast Cancer Cell Lines. Molecular Cancer Research 2007, 5 (2), 195–201. 10.1158/1541-7786.MCR-06-0263.

(32) Falvo, E.; Tremante, E.; Fraioli, R.; Leonetti, C.; Zamparelli, C.; Boffi, A.; Morea, V.; Ceci, P.; Giacomini, P. Antibody–Drug Conjugates: Targeting Melanoma with Cisplatin Encapsulated in Protein-Cage Nanoparticles Based on Human Ferritin. Nanoscale 2013, 5 (24), 12278. 10.1039/c3nr04268e.

(33) Bellini, M.; Mazzucchelli, S.; Galbiati, E.; Sommaruga, S.; Fiandra, L.; Truffi, M.; Rizzuto, M. A.; Colombo, M.; Tortora, P.; Corsi, F.; Prosperi, D. Protein Nanocages for Self-Triggered Nuclear Delivery of DNA-Targeted Chemotherapeutics in Cancer Cells. J Control Release 2014, 196, 184–196. 10.1016/j.jconrel.2014.10.002.

(34) Kappel, C.; Seidl, C.; Medina-Montano, C.; Schinnerer, M.; Alberg, I.; Leps, C.; Sohl, J.; Hartmann, A.-K.; Fichter, M.; Kuske, M.; Schunke, J.; Kuhn, G.; Tubbe, I.; Paßlick, D.; Hobernik, D.; Bent, R.; Haas, K.; Montermann, E.; Walzer, K.; Diken, M.; Schmidt, M.; Zentel, R.; Nuhn, L.; Schild, H.; Tenzer, S.; Mailänder, V.; Barz, M.; Bros, M.; Grabbe, S. Density of Conjugated Antibody Determines the Extent of Fc Receptor Dependent Capture of Nanoparticles by Liver Sinusoidal Endothelial Cells. ACS Nano 2021, 15 (9), 15191–15209. 10.1021/acsnano.1c05713.

(35) Metkar, S. S.; Wang, B.; Ebbs, M. L.; Kim, J. H.; Lee, Y. J.; Raja, S. M.; Froelich, C. J. Granzyme B Activates Procaspase-3 Which Signals a Mitochondrial Amplification Loop for Maximal Apoptosis. J Cell Biol 2003, 160 (6), 875–885. 10.1083/jcb.200210158.

(36) Cao, H.; Schroeder, B.; Chen, J.; Schott, M. B.; McNiven, M. A. The Endocytic Fate of the Transferrin Receptor Is Regulated by C-Abl Kinase. J Biol Chem 2016, 291 (32), 16424–16437. 10.1074/jbc.M116.724997.

(37) Mayle, K. M.; Le, A. M.; Kamei, D. T. The Intracellular Trafficking Pathway of Transferrin. Biochim Biophys Acta 2012, 1820 (3), 264–281. 10.1016/j.bbagen.2011.09.009.

(38) Sigismund, S.; Argenzio, E.; Tosoni, D.; Cavallaro, E.; Polo, S.; Di Fiore, P. P. Clathrin-Mediated Internalization Is Essential for Sustained EGFR Signaling but Dispensable for Degradation. Developmental Cell 2008, 15 (2), 209–219. 10.1016/j.devcel.2008.06.012.

(39) Henriksen, L.; Grandal, M. V.; Knudsen, S. L. J.; van Deurs, B.; Grøvdal, L. M. Internalization Mechanisms of the Epidermal Growth Factor Receptor after Activation with Different Ligands. PLoS One 2013, 8 (3), e58148. 10.1371/journal.pone.0058148.

(40) Mazzucchelli, S.; Bellini, M.; Fiandra, L.; Truffi, M.; Rizzuto, M. A.; Sorrentino, L.; Longhi, E.; Nebuloni, M.; Prosperi, D.; Corsi, F. Nanometronomic Treatment of 4T1 Breast Cancer with Nanocaged Doxorubicin Prevents Drug Resistance and Circumvents Cardiotoxicity. Oncotarget 2017, 8 (5), 8383–8396. 10.18632/oncotarget.14204.

(41) Sevieri, M.; Sitia, L.; Bonizzi, A.; Truffi, M.; Mazzucchelli, S.; Corsi, F. Tumor Accumulation and Off-Target Biodistribution of an Indocyanine-Green Fluorescent Nanotracer: An Ex Vivo Study on an Orthotopic Murine Model of Breast Cancer. IJMS 2021, 22 (4), 1601. 10.3390/ijms22041601.

(42) Sevieri, M.; Pinori, M.; Chesi, A.; Bonizzi, A.; Sitia, L.; Truffi, M.; Morasso, C.; Corsi, F.; Mazzucchelli, S. Novel Bioengineering Strategies to Improve Bioavailability and In Vivo Circulation of H-Ferritin Nanocages by Surface Functionalization. ACS Omega 2023, 8 (8), 7244–7251. 10.1021/acsomega.2c07794.

(43) Falvo, E.; Malagrinò, F.; Arcovito, A.; Fazi, F.; Colotti, G.; Tremante, E.; Di Micco, P.; Braca, A.; Opri, R.; Giuffrè, A.; Fracasso, G.; Ceci, P. The Presence of Glutamate Residues on the PAS Sequence of the Stimuli-Sensitive Nano-Ferritin Improves *in Vivo* Biodistribution and Mitoxantrone Encapsulation Homogeneity. Journal of Controlled Release 2018, 275, 177–185. 10.1016/j.jconrel.2018.02.025.

(44) Pinilla-Macua, I.; Grassart, A.; Duvvuri, U.; Watkins, S. C.; Sorkin, A. EGF Receptor Signaling, Phosphorylation, Ubiquitylation and Endocytosis in Tumors in Vivo. Elife 2017, 6, e31993. 10.7554/eLife.31993.

(45) Bitsikas, V.; Corrêa, I. R.; Nichols, B. J. Clathrin-Independent Pathways Do Not Contribute Significantly to Endocytic Flux. Elife 2014, 3, e03970. 10.7554/eLife.03970.

(46) Tan, X.; Lambert, P. F.; Rapraeger, A. C.; Anderson, R. A. Stress-Induced EGFR Trafficking: Mechanisms, Functions, and Therapeutic Implications. Trends Cell Biol 2016, 26 (5), 352–366. 10.1016/j.tcb.2015.12.006.

(47) Rennick, J. J.; Johnston, A. P. R.; Parton, R. G. Key Principles and Methods for Studying the Endocytosis of Biological and Nanoparticle Therapeutics. Nat. Nanotechnol. 2021, 16 (3), 266–276. 10.1038/s41565-021-00858-8.

(48) Matthaeus, C.; Taraska, J. W. Energy and Dynamics of Caveolae Trafficking. Front. Cell Dev. Biol. 2021, 8. 10.3389/fcell.2020.614472.

(49) Innocenti, M.; Gerboth, S.; Rottner, K.; Lai, F. P. L.; Hertzog, M.; Stradal, T. E. B.; Frittoli, E.; Didry, D.; Polo, S.; Disanza, A.; Benesch, S.; Fiore, P. P. D.; Carlier, M.-F.; Scita, G. Abi1 Regulates the Activity of N-WASP and WAVE in Distinct Actin-Based Processes. Nat Cell Biol 2005, 7 (10), 969–976. 10.1038/ncb1304.

(50) Galovic, M.; Xu, D.; Areces, L. B.; van der Kammen, R.; Innocenti, M. Interplay between N-WASP and CK2 Optimizes Clathrin-Mediated Endocytosis of EGFR. J Cell Sci 2011, 124 (Pt 12), 2001–2012. 10.1242/jcs.081182.

(51) Isogai, T.; van der Kammen, R.; Leyton-Puig, D.; Kedziora, K. M.; Jalink, K.; Innocenti, M. Initiation of Lamellipodia and Ruffles Involves Cooperation between mDia1 and the Arp2/3 Complex. J Cell Sci 2015, 128 (20), 3796–3810. 10.1242/jcs.176768.

(52) Wilkins, M. R.; Gasteiger, E.; Bairoch, A.; Sanchez, J. C.; Williams, K. L.; Appel, R. D.; Hochstrasser, D. F. Protein Identification and Analysis Tools in the ExPASy Server. Methods Mol Biol 1999, 112, 531–552. 10.1385/1-59259-584-7:531.

(53) Isogai, T.; van der Kammen, R.; Bleijerveld, O. B.; Goerdayal, S. S.; Argenzio, E.; Altelaar, A. F.; Innocenti, M. Quantitative Proteomics Illuminates a Functional Interaction between mDia2 and the Proteasome. J Proteome Res 2016, 15 (12), 4624–4637. 10.1021/acs.jproteome.6b00718.

(54) Leyton-Puig, D.; Isogai, T.; Argenzio, E.; van den Broek, B.; Klarenbeek, J.; Janssen, H.; Jalink, K.; Innocenti, M. Flat Clathrin Lattices Are Dynamic Actin-Controlled Hubs for Clathrin-Mediated Endocytosis and Signalling of Specific Receptors. Nature communications 2017, 8, 16068. 10.1038/ncomms16068.

(55) Leyton-Puig, D.; Kedziora, K. M.; Isogai, T.; van den Broek, B.; Jalink, K.; Innocenti, M. PFA Fixation Enables Artifact-Free Super-Resolution Imaging of the Actin Cytoskeleton and Associated Proteins. Biology open 2016, 5 (7), 1001– 1009. 10.1242/bio.019570.

(56) Dunn, K. W.; Kamocka, M. M.; McDonald, J. H. A Practical Guide to Evaluating Colocalization in Biological Microscopy. Am J Physiol Cell Physiol 2011, 300 (4), C723–C742. 10.1152/ajpcell.00462.2010.

(57) Carpenter, A. E.; Jones, T. R.; Lamprecht, M. R.; Clarke, C.; Kang, I. H.; Friman, O.; Guertin, D. A.; Chang, J. H.; Lindquist, R. A.; Moffat, J.; Golland, P.; Sabatini, D. M. CellProfiler: Image Analysis Software for Identifying and Quantifying Cell Phenotypes. Genome Biology 2006, 7 (10), R100. 10.1186/gb-2006-7-10-r100.

(58) Argenzio, E.; Innocenti, M. The Chloride Intracellular Channel Protein CLIC4 Inhibits Filopodium Formation Induced by Constitutively Active Mutants of Formin mDia2. FEBS Lett 2020, 594 (11), 1750–1758. 10.1002/1873-3468.13766.

