## Supplemental figures for "Dual-targeting strategy to repurpose Cetuximab with HFn nanoconjugates for immunotherapy of triple-negative breast cancer"

### Supplementary Tables

**Supplementary table S1. Physicochemical properties of HFn-CTX nanoconjugates.**

| Reaction | Purity | HFn yield | CTX yield | Dimensions |
|---|---|---|---|---|
| HFn-CTX | 77.5 % | 31.7 ± 4.81 % | 31.7 ± 4.82 % | 28.2 ± 4.15 nm |

Values are the mean of three different batches of HFn-CTX.

**Supplementary table S2. Optimization of HFn labeling with AF488.**

| Reaction | HFn:AF488 (molar ratio) | Protein concentration (mg mL$^{-1}$) | Reaction conditions | DOL |
|---|---|---|---|---|
| 1 | 1:2.5 | 2 | 2 h RT | 0.15 |
| 2 | 1:6 | 2 | 2 h RT | 0.3 |
| 3 | 1:30 | 2 | 30 min RT then o/n 4 °C | 1.5 |
| 4 | 1:50 | 5 | 30 min RT then o/n 4°C | 3.9 |
| 5 | 1:100 | 2 | 30 min RT then o/n 4 °C | 4 |

Summary of the different conditions tested.

**Supplementary table S3. Optimization of CTX labeling with AF647.**

| Reaction | Molar ratio CTX:AF647 | Protein concentration (mg mL$^{-1}$) | Reaction conditions | PEG presence? | DOL |
|---|---|---|---|---|---|
| 1 | 1:8.9 | 2 | 1 h RT | 1:40 | 1.1 |
| 2 | 1:20 | 2 | 1 h RT | 1:40 | 2.1 |
| 3 | 1:10 | 5 | 2.5 h RT | 1:40 | 4.6 |
| 4 | 1:15 | 5 | 2.5 h RT | 1:40 | 7.9 |
| 4a | 1:15 | 5 | 2.5 h RT | No PEG | 9.1 |

Summary of the different conditions tested.

**Supplementary table 4. Optimization of HFn labelling with AF647.**

| Reaction | HFn:AF647 (molar ratio) | Protein concentration (mg $mL^{-1}$) | Reaction conditions | DOL |
|---|---|---|---|---|
| 1 | 1:25 | 5 | 30 min RT then o/n 4 °C | 3.9 |
| 2 | 1:50 | 5 | 30 min RT then o/n 4 °C | 6.04 |

Due to the high tissue autofluorescence at wavelengths across 488 nm, we did not use AF488 for the in vivo experiments and decided to label HFn with AF647. Based on the results shown in Supplementary table 2, we used a molar ratio of 1:25, which gave an acceptable degree of labelling while saving fluorophore. For CTX-AF750, we used a molar ratio of 1:10.

**Supplementary table 5. Optimization of CTX labelling with AF750.**

| Reaction | Molar ratio CTX:AF750 | Protein concentration (mg $mL^{-1}$) | Reaction conditions | Molar ratio CTX:PEG | DOL |
|---|---|---|---|---|---|
| 1 | 1:10 | 5 | 2.5 h RT | 1:40 | 4.9 |
| 2 | 1:5 | 5 | 2.5 h RT | No PEG | 3.5 |
| 3 | 1:2 | 5 | 2.5 h RT | No PEG | 0.8 |

Due to the high tissue autofluorescence at wavelengths across 488 nm, we did not use AF488 for the in vivo experiments. and decided to label CTX with AF750. Based on the results shown in Supplementary table 3, we used a molar ratio of 1:10.

A)

B)

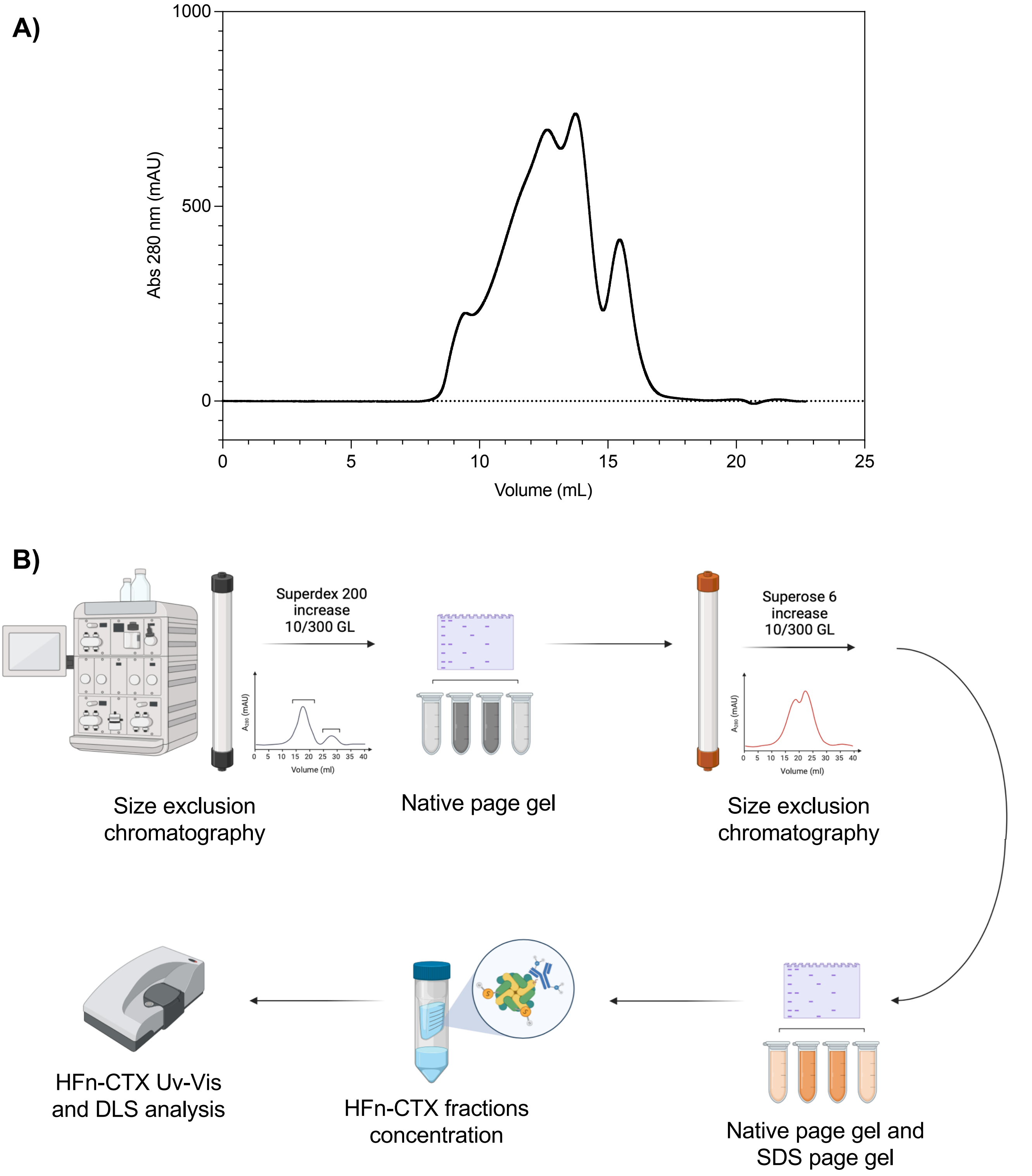
1000
500
0
Abs 280 nm (mAU)
0
5
10
15
20
25
Volume (mL)
Superdex 200
increase
10/300 GL
Volume (ml)
Size exclusion
chromatography
Native page gel
Superose 6
increase
10/300 GL
Volume (ml)
Size exclusion
chromatography
HFn-CTX Uv-Vis
and DLS analysis
HFn-CTX fractions
concentration
Native page gel and
SDS page gel

Figure S1

**Figure S1. Two-step purification of the HFn-CTX nanoconjugate by SEC-FPLC.**

**(A)** Representative elution profile of the HFn-CTX nanoconjugate on Superose 6 column. **(B)** Schematic representation of the steps required to purify the HFn-CTX nanoconjugate. After the conjugation reaction, the nanoconjugate was collected and purified through two SEC-FPLC steps, the former and the latter employing a Superdex 200 increase 10/300 GL column and a Superose 6 increase 10/300 GL column equilibrated with PBS, respectively. The eluted fractions were then characterized by SDS-PAGE, Western blot, and dynamic light scattering. The fractions containing the HFn-CTX nanoconjugate were concentrated on Amicon filters (100 kDa MWCO), if deemed necessary. Concentrated HFn-CTX nanoconjugate was aliquoted and kept at 4 °C in PBS after assessing its stability by means of UV-VIS spectrometry and/or centrifugation.

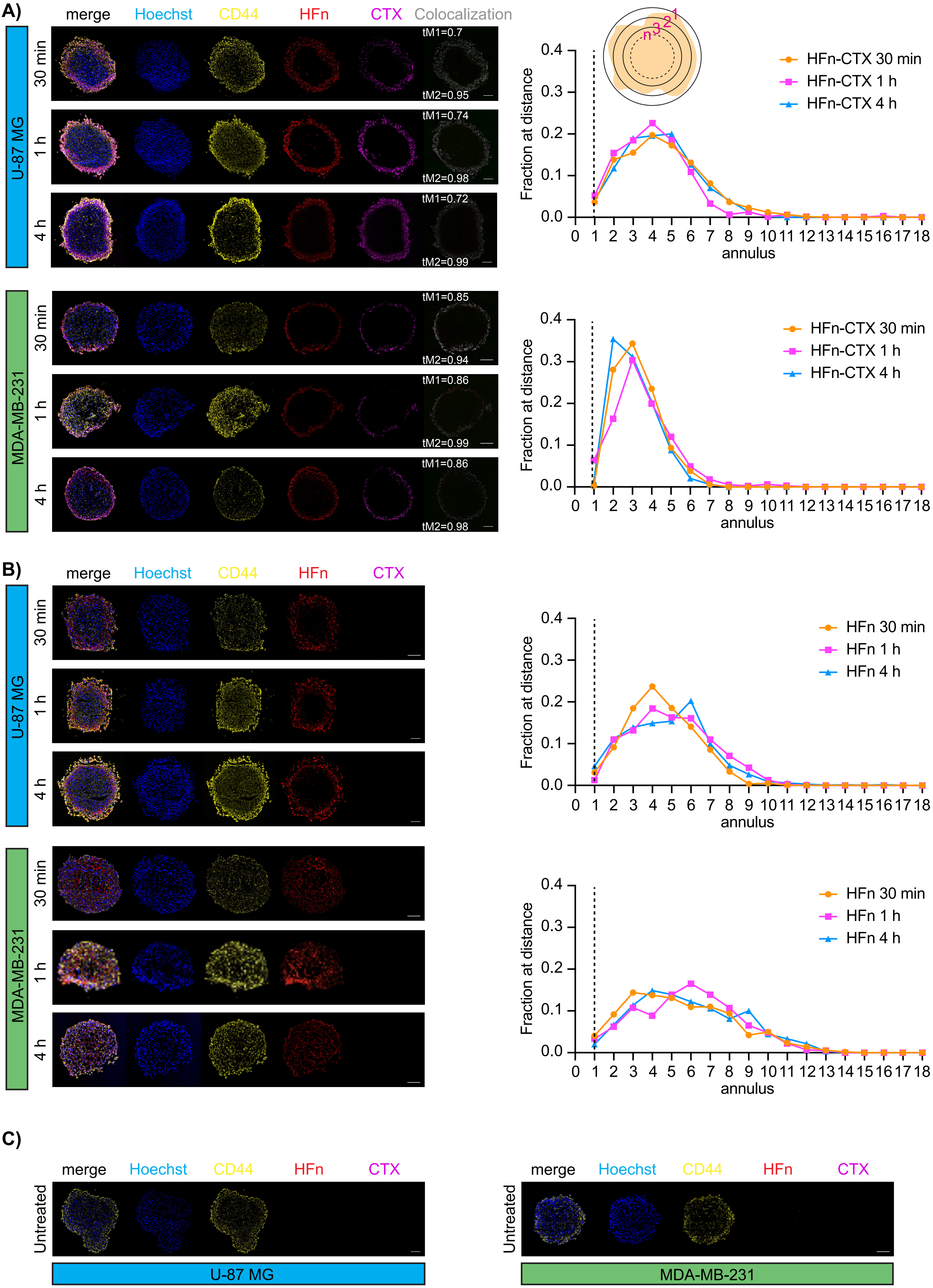
A)
merge
Hoechst
CD44
HFn
CTX
Colocalization
U-87 MG
30 min
1 h
4 h
tM1=0.7
tM2=0.95
tM1=0.74
tM2=0.98
tM1=0.72
tM2=0.99
MDA-MB-231
tM1=0.85
tM2=0.94
tM1=0.86
tM2=0.99
tM1=0.86
tM2=0.98
Fraction at distance
annulus
HFn-CTX 30 min
HFn-CTX 1 h
HFn-CTX 4 h
B)
HFn 30 min
HFn 1 h
HFn 4 h
C)
Untreated
U-87 MG
MDA-MB-231

Figure S2

**Figure S2. Distribution of the HFn-CTX nanoconjugate and HFn nanoparticles within GBM and TNBC spheroids.**

**(A-C)** *(Left)* Representative cryosections of U-87 MG and MDA-MB-231 spheroids incubated with double labelled HFn-CTX **(A)**, or AF647-labelled HFn **(B)** (0.1 mg $mL^{-1}$ each) for 30 min, 1 and 4 hours, or left untreated as a control **(C)**. Plasma membrane stained with CD44 is depicted in yellow, nuclei stained with Hoechst in blue, HFn-AF647 (HFn) in red, CTX-AF750 (CTX) in magenta. Colocalization maps (Colocalization) are shown in gray. Mander's coefficient tM1 (CTX-AF750 signal *vs.* HFn-AF647 signal) and tM2 (HFn-AF647 signal *vs.* CTX-AF750) are also indicated. Scale bar, 100 µm. **(A,B)** *(Right)* Automated analysis of the distribution of the HFn-CTX nanoconjugate and HFn nanoparticles in U-87 MG and MDA-MB-231 spheroids. Spheroids were segmented in 10-pixel wide round annuli and the fraction of the total HFn-AF647 signal within each annulus (Fraction at a distance) calculated with CellProfiler (see schematic on top). Each image was segmented in 19 annuli centered on the spheroid's center. Given that the radius was variable, the annulus containing the spheroid's border was set to 1. This registration allowed a precise comparison of the signal distribution among the analysed time points (30 minutes in ochre, 1 hour in magenta, 4 hours in cyan).

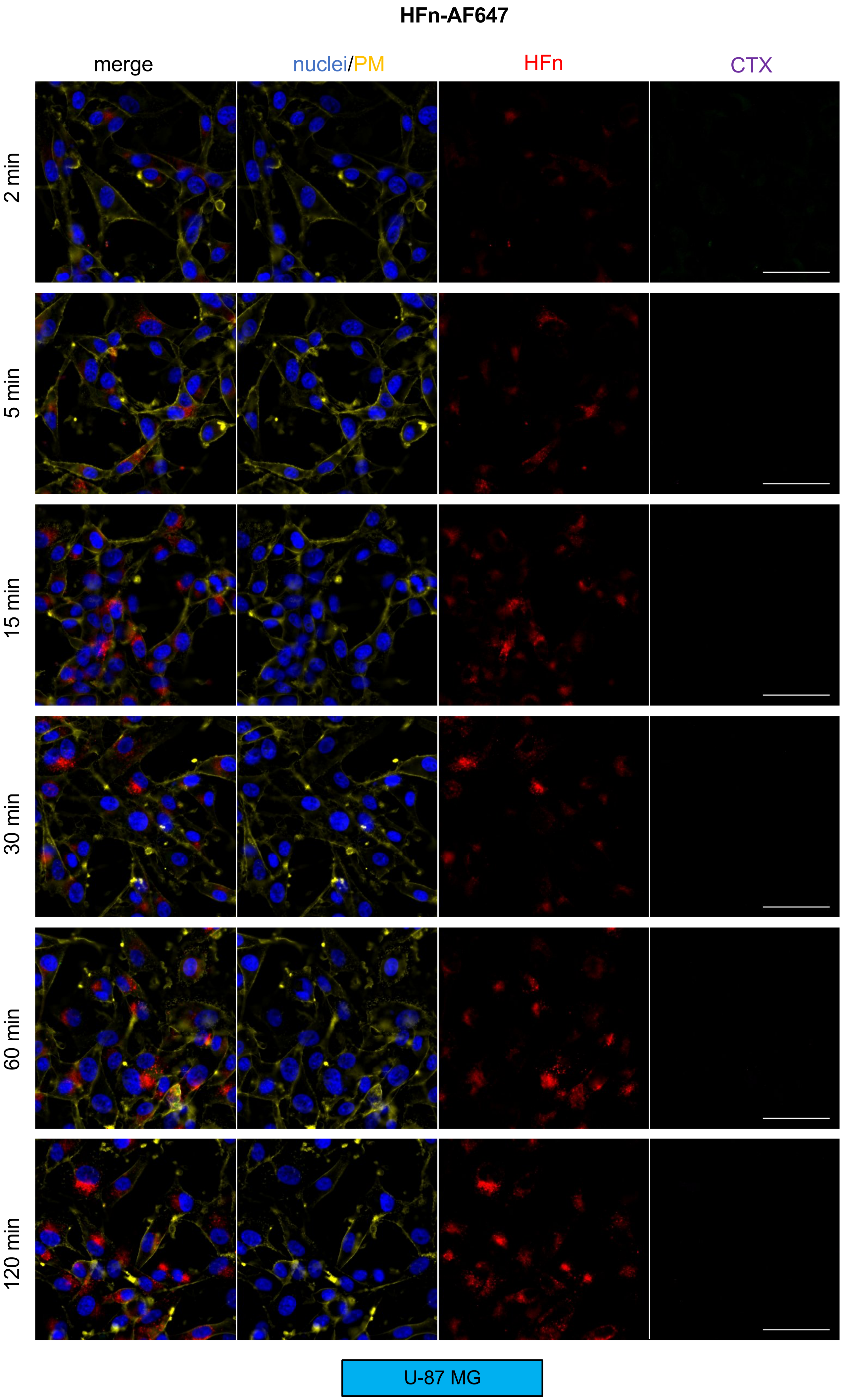
HFn-AF647
merge
nuclei/PM
HFn
CTX
2 min
5 min
15 min
30 min
60 min
120 min
U-87 MG

Figure S3

### HFn-AF647-CTX-AF750

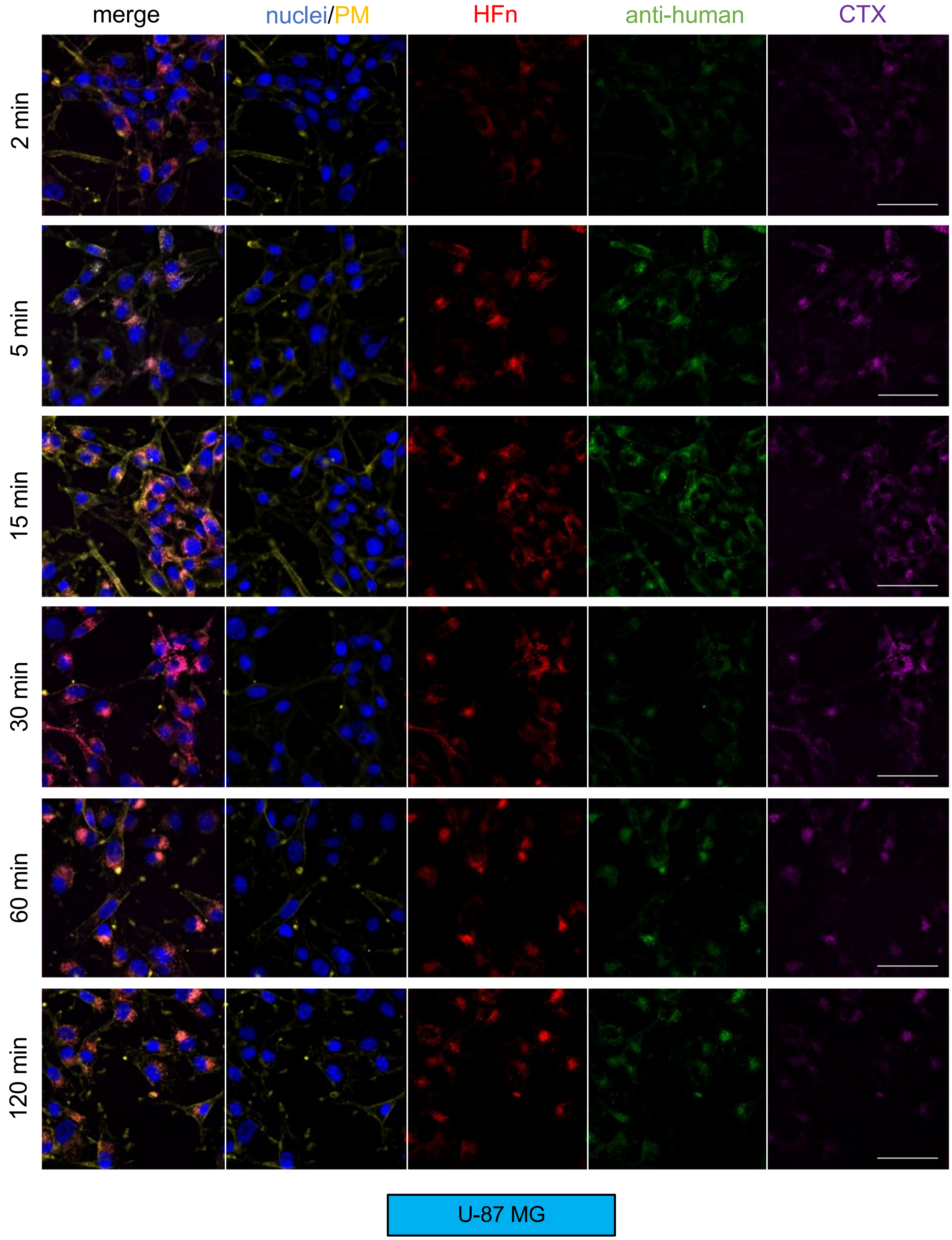


Figure S3

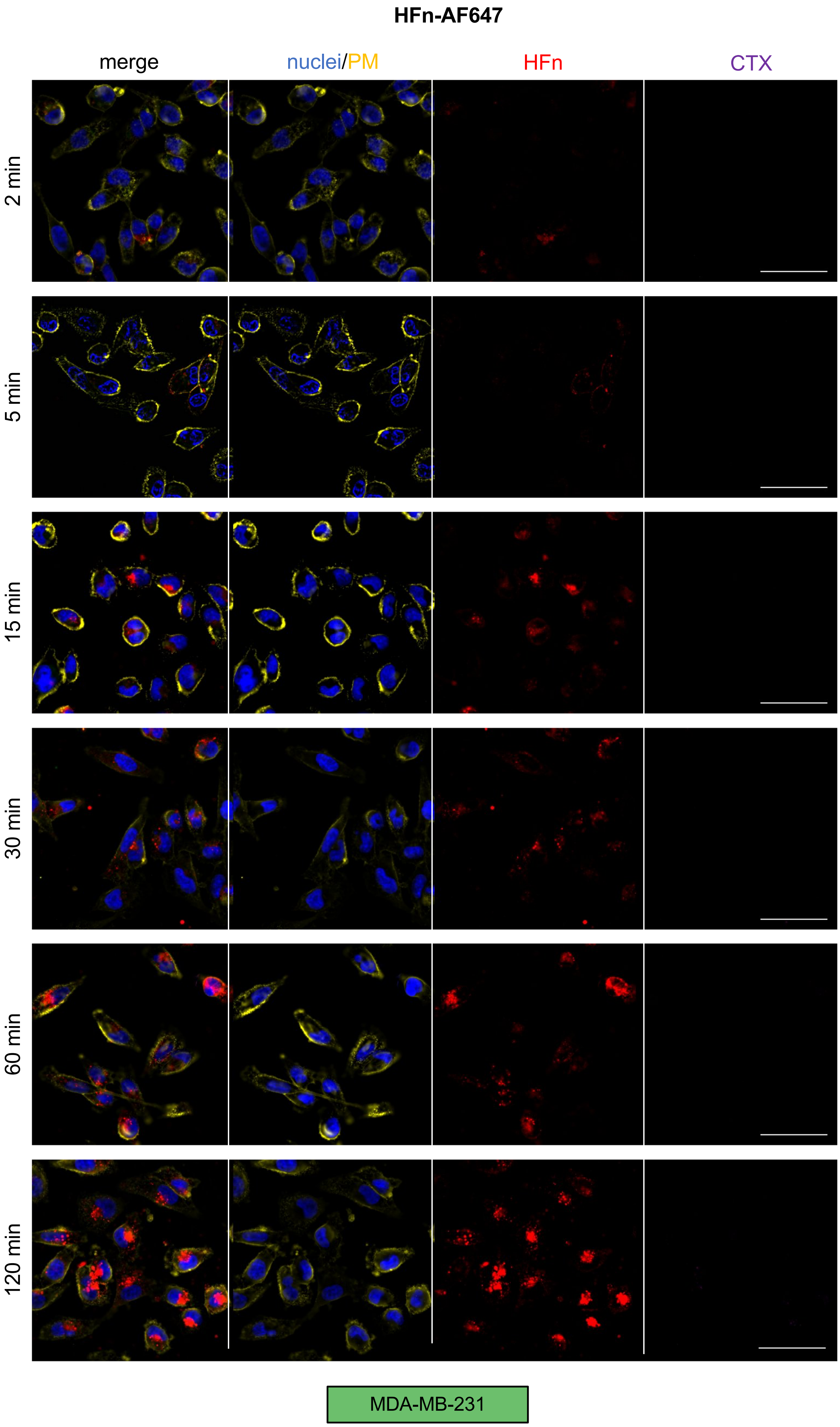
HFn-AF647
merge
nuclei/PM
HFn
CTX
2 min
5 min
15 min
30 min
60 min
120 min
MDA-MB-231

Figure S3

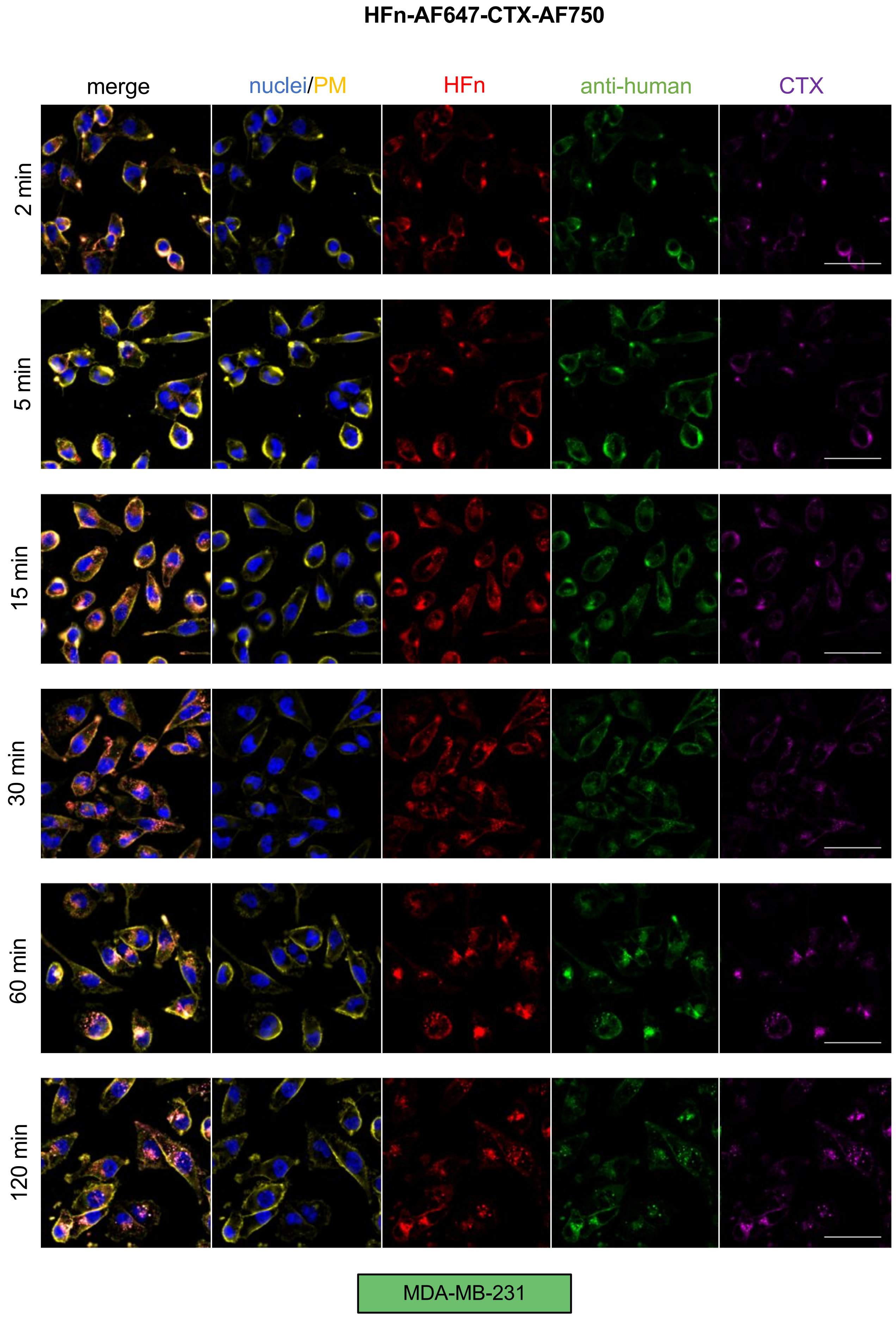
HFn-AF647-CTX-AF750
merge
nuclei/PM
HFn
anti-human
CTX
2 min
5 min
15 min
30 min
60 min
120 min
MDA-MB-231

Figure S3

**Figure S3. Large-field images of the uptake of the HFn-CTX nanoconjugate or HFn nanoparticles in U-87 MG and MDA-MB-231 cells**

Representative images of U-87 MG and MDA-MB-231 cells incubated with double labelled HFn-CTX or labelled HFn (0.1 mg $mL^{-1}$ each) for 2, 5, 25, 30, 60 and 120 min. Samples were processed as in Figure 3 and Figure 4. Zooms of U-87 MG and MDA-MB-231 cells are depicted in Figure 3 and Figure 4, respectively. Scale bar, 100 µm.

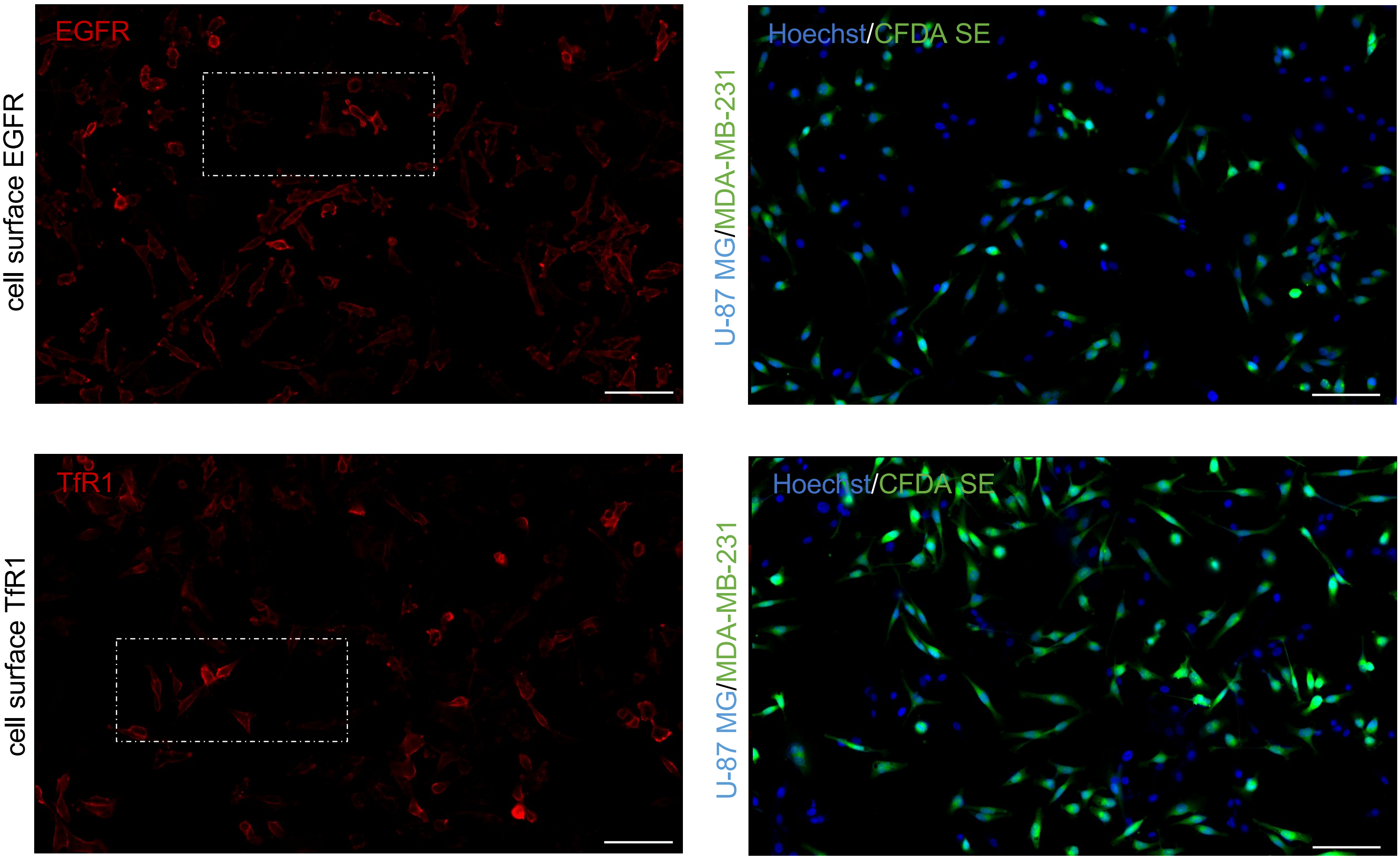
EGFR
cell surface EGFR
Hoechst/CFDA SE
U-87 MG/MDA-MB-231
TfR1
cell surface TfR1
Hoechst/CFDA SE
U-87 MG/MDA-MB-231

Figure S4

**Figure S4. Large-field images of cell-surface EGFR and TfR1 levels in U-87 MG and MDA-MB-231 co-cultures.**

Samples were processed as in Figure 5D: EGFR (top) and TfR1 (bottom) are depicted in red, nuclei in blue, and MDA-MB-231 are in green. The white rectangles represent the area shown in Figure 5D. Scale bar 100 µm.

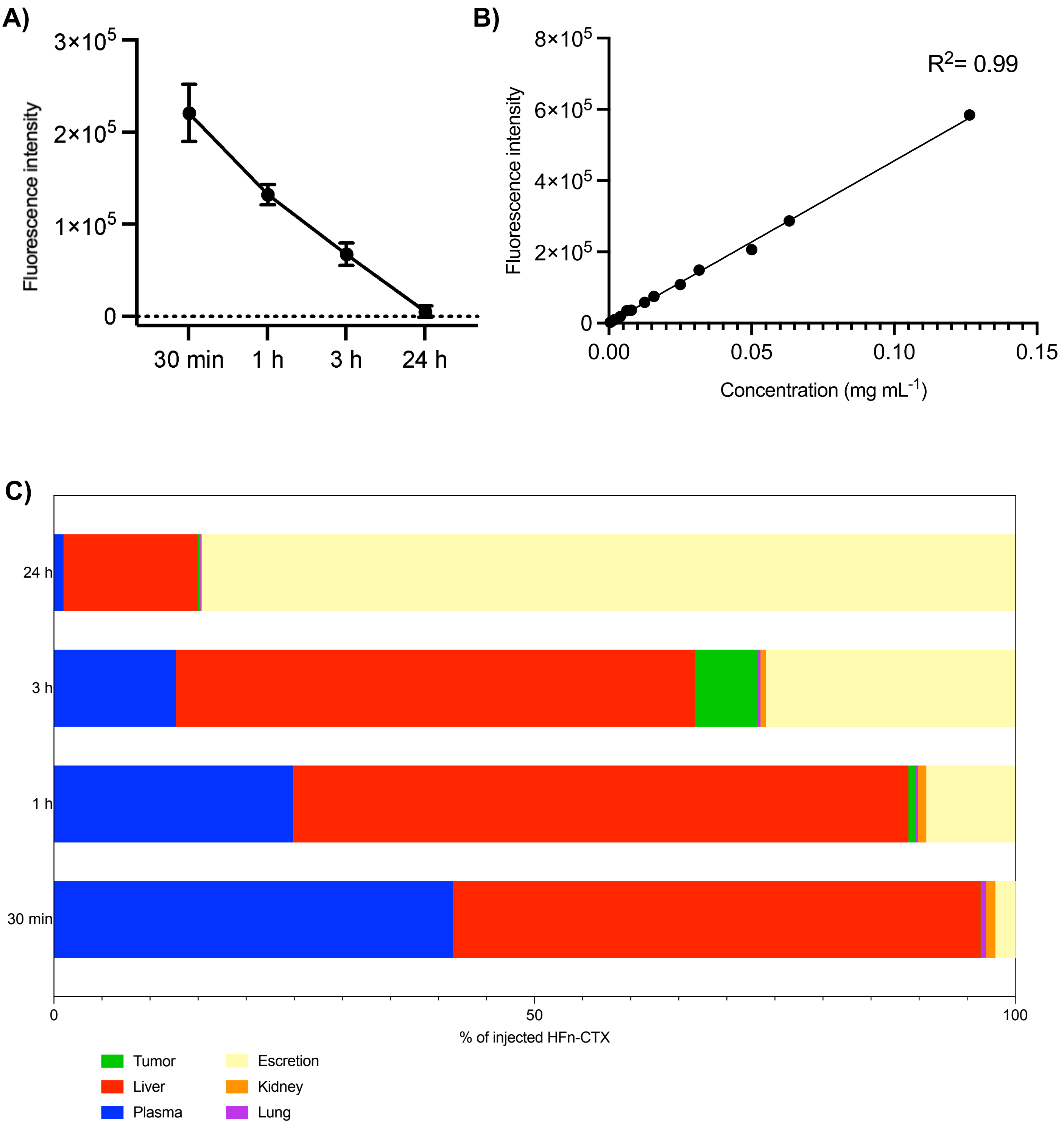
A)
3×10^5
2×10^5
1×10^5
0
Fluorescence intensity
30 min
1 h
3 h
24 h
B)
8×10^5
6×10^5
4×10^5
2×10^5
0
Fluorescence intensity
R^2= 0.99
0.00
0.05
0.10
0.15
Concentration (mg mL^-1)
C)
24 h
3 h
1 h
30 min
0
50
100
% of injected HFn-CTX
Tumor
Liver
Plasma
Escretion
Kidney
Lung

Figure S5

**Figure S5**. **Biodistribution of the HFn-CTX nanoconjugate in mice.**

**(A)** Fluorescence intensity of the HFn-CTX nanoconjugate in mouse plasma at the indicated time points. **(B)** Calibration curve used to determine the percentage of the HFn-CTX nanoconjugate retained in plasma. **(C)** Percentage of the injected dose (ID) found in tumors, off-target organs, and plasma. The percentage of excreted HFn-CTX was calculated by subtraction.
